# CRISPR screens reveal ZBTB17/MIZ1 as a peroxisome regulator

**DOI:** 10.1101/2024.07.25.605214

**Authors:** Hongqin Liu, Xi Chen, Hanlin Wang, Guanglei Zhuang, Zheng-Jiang Zhu, Min Zhuang

**Affiliations:** School of Life Science and Technology, ShanghaiTech University, Shanghai, 201210, China; University of Chinese Academy of Sciences, Beijing, 100049, China; Interdisciplinary Research Center on Biology and Chemistry, Shanghai Institute of Organic Chemistry, Chinese Academy of Sciences, Shanghai, 200032 China; State Key Laboratory of Chemical Biology, Shanghai Institute of Materia Medica, Chinese Academy of Sciences, Shanghai, 201203, China; Shandong Laboratory of Yantai Drug Discovery, Bohai Rim Advanced Research Institute for Drug Discovery, Yantai, Shandong 264117, China; State Key Laboratory of Systems Medicine for Cancer, Department of Obstetrics and Gynecology, Shanghai Cancer Institute, Ren Ji Hospital, Shanghai Jiao Tong University School of Medicine, Shanghai, China; Shanghai Key Laboratory of Gynecologic Oncology, Ren Ji Hospital, Shanghai Jiao Tong University School of Medicine, Shanghai, China; Shanghai Key Laboratory of Aging Studies, Shanghai, 201210 China

## Abstract

Peroxisomes are integral metabolic organelles involved in both catabolic and anabolic processes in humans, with defects often linked to diseases. The functions of peroxisomes are regulated at transcriptional, translational, and post-translational levels. In this study, we employed the CRISPR/Cas9-based genetic screening of a ubiquitin ligase library to identify regulators of human peroxisomes. We discovered that ZBTB17 (also referred as MIZ1) plays a role in regulating the import of proteins into peroxisomes. Independent of its ubiquitin ligase activity, ZBTB17/MIZ1 operates as a transcription factor to directly modulate the expression of key importer PEX13, thereby influencing the localization of peroxisomal enzymes. Furthermore, metabolomic profiling reveals that the knockdown of *ZBTB17* or *PEX13* results in similar metabolic alterations, characterized by downregulated purine synthesis, suggesting that ZBTB17’s role in metabolic regulation likely operates through peroxisomes. Collectively, we identify ZBTB17 as a key regulator of peroxisomal protein import, thereby affecting peroxisomal function and nucleotide metabolism. Our findings provide insights into the multifaceted regulation of peroxisomes in complex human cells and shed light on the molecular mechanisms underlying ZBTB17’s role as a transcriptional regulator.

## INTRODUCTION

Peroxisomes are dynamic, membrane-bound organelles that play crucial roles in numerous cellular metabolic reactions, including fatty acid β-oxidation, ether lipid and bile acid synthesis, cholesterol transport, reactive oxygen species (ROS) metabolism, and anti-viral signaling (Chu et al., 2015; Dixit et al., 2010; Islinger et al., 2018; Smith and Aitchison, 2013). The size and quantity of peroxisomes can greatly vary among different species. Mammalian cells maintain peroxisome homeostasis through a balance between regulated biogenesis and degradation (Mahalingam et al., 2021). Many PEX genes have been characterized, encoding peroxin proteins that operate in various stages of peroxisome biogenesis. These include PEX3, PEX19, and PEX16 for membrane formation (Hohfeld et al., 1991; Kim and Mullen, 2013; Matsuzaki and Fujiki, 2008; Shibata et al., 2004), PEX1, 2, 5, 6, 7, 10, 12, 13, 14 for matrix protein import (Dammai and Subramani, 2001; El Magraoui et al., 2012; Feng et al., 2022; Meinecke et al., 2010; Platta et al., 2007; Titorenko and Rachubinski, 2000; Walton et al., 1995), and PEX11 for peroxisome proliferation (Chang et al., 2015; Thomas et al., 2015). Mutations in *PEX* genes can result in peroxisome biogenesis disorders, whereby patients have reduced or no peroxisomes and often face early mortality (Francis et al., 1995; Gould and Valle, 2000; Zalckvar and Schuldiner, 2022). Dysfunctions in peroxisomes have also been associated with neurodegenerative disease, aging, cancer, and type 2 diabetes (Waterham et al., 2016; Zalckvar and Schuldiner, 2022).

Peroxisome homeostasis is highly regulated at different levels. Notably, the biogenesis of peroxisomes is known to be controlled at the transcriptional level. In the methylotrophic yeast *Pichia pastoris*, the transcription factor Mxr1p induces the expression of several *PEX* genes and metabolic enzymes (Lin-Cereghino et al., 2006). Similarly, in human cells, the PPAR (peroxisome proliferator-activated receptor) transcription factor family responds to lipid activators, initiating the biogenesis and proliferation of peroxisomes (Lemberger et al., 1996; Rangwala and Lazar, 2004; Zhang et al., 2013). The biogenesis and degradation of peroxisomes are also tightly regulated by post-translational modifications, with ubiquitination playing a significant role (Platta et al., 2007; Sargent et al., 2016; Zheng et al., 2022). Specifically, the PEX2/10/12 ubiquitin ligase complex is necessary for matrix protein import during peroxisome biogenesis (El Magraoui et al., 2012; Feng et al., 2022). In contrast, overexpression of PEX2 triggers pexophagy (Sargent et al., 2016), a process leading to the programmed degradation of peroxisomes *via* autophagy. In addition, amino acid depletion or mTOR inhibition also induces pexophagy, but in a manner dependent on the ubiquitin ligase MARCH5 (Zheng et al., 2022). Therefore, ubiquitin ligases are instrumental in regulating both peroxisome biogenesis and degradation.

Genetic screening in yeast has identified numerous *PEX* genes important for peroxisome biogenesis (Distel et al., 1996), and these genes exhibit a high degree of conservation in mammalian cells. However, given the larger number of peroxisomes in human cells compared to the limited number typically found in yeast, it is expected that the regulation of peroxisomes in human cells is more complex. This complexity could extend to both the organelle level, through the regulation of peroxisome numbers, and the enzyme level, by controlling the import of peroxisomal metabolic enzymes. Such regulatory mechanisms would allow the fine-tuning of peroxisome functions. In this study, we hypothesized that there might be other peroxisome regulators in human cells. To investigate this, we employed an sgRNA library targeting ubiquitin ligases to screen for potential peroxisome regulators using CRISPR/Cas9 technology. We identified and confirmed the regulatory role of ZBTB17 in peroxisome protein import. ZBTB17 possesses a domain architecture comprising both the BTB domain, which is essential for ubiquitin ligase activity, and zinc finger motifs, which are necessary for DNA binding in transcriptional regulation. Further characterization revealed ZBTB17 as a transcription factor that directly controls *PEX13* expression to regulate peroxisomal protein import, thereby influencing cellular metabolism.

## RESULTS

### Identification of potential peroxisome regulators by the CRISPR/Cas9 screen

To perform genetic screen in HeLa cells, we first generated a HeLa cell line stably expressing spCas9 (HeLa-Cas9). The expression of spCas9 in various clones was confirmed using anti-Flag immunoblots, and its activity was validated using the T7 assay (**Fig. S1A** and **1B**). To generate a peroxisome reporter, we design a construct containing both GFP-SKL (poGFP) and mCherry with an IRES (internal ribosome entry site) sequence in between (**Fig. 1A**). SKL is a peroxisome-targeting peptide composed of three amino acids Ser-Lys-Leu. GFP-SKL mainly localizes in peroxisomes while mCherry remains in the cytoplasm. The GFP/mCherry ratio can be used to monitor the change of peroxisomes in cells. We generated a stable cell line expressing GFP(SKL)-IRES-mCherry on the background of HeLa-Cas9 cells. The expression of both GFP-SKL and mCherry were confirmed with flow cytometry (**Fig. S1C**) and we name this cell line as HeLa-Cas9-poGFP/mCherry.

**Figure 1.**
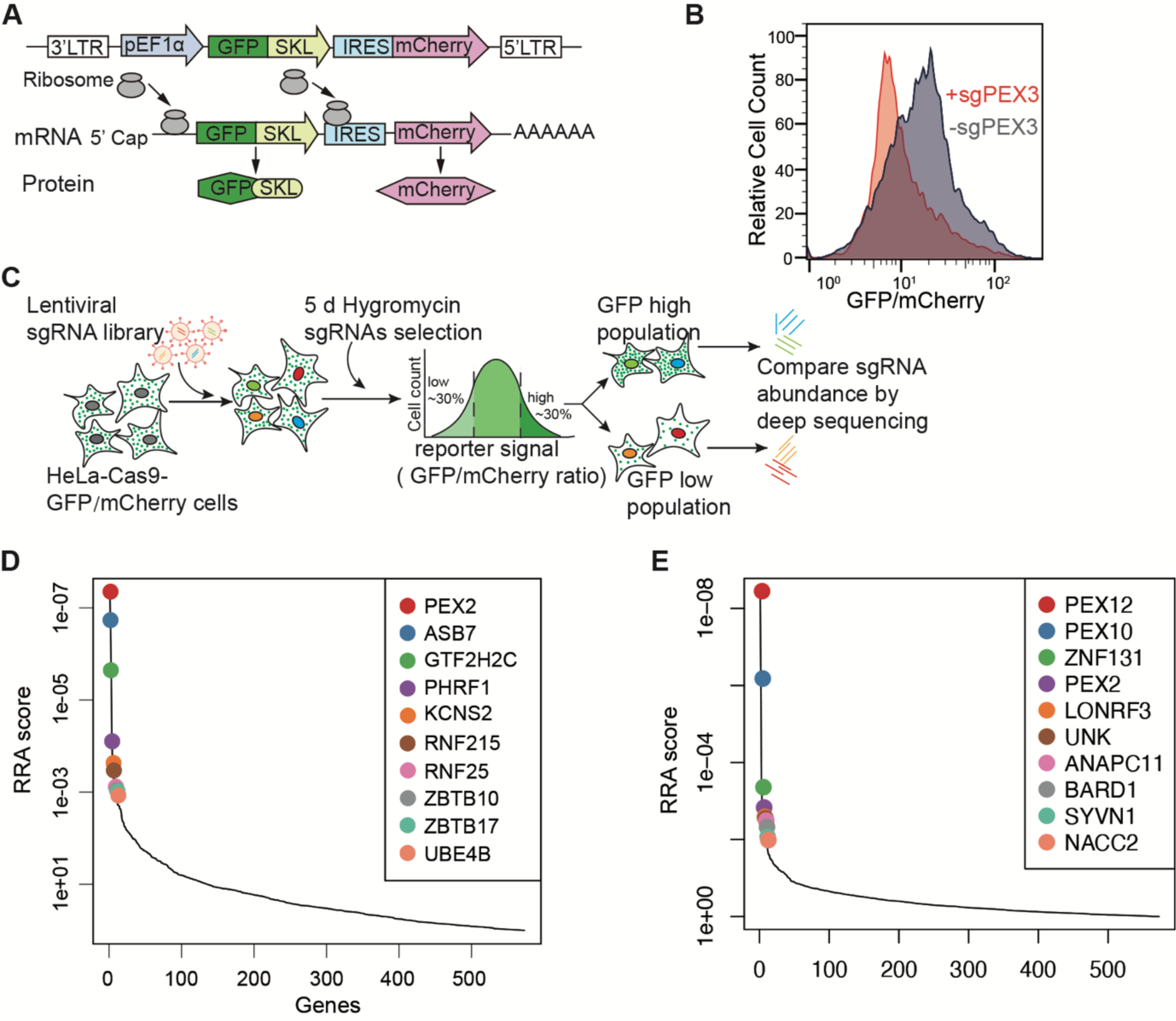
Identification of peroxisome regulators using a dual-fluorescence reporter system coupled with a CRISPR screen. **(A)** Diagram of the peroxisome reporter pEF1α-GFP(SKL)-IRES-mCherry. SKL stands for Ser-Lys-Leu fused to the C terminus of GFP; IRES stands for internal ribosome entry site. **(B)** Validation of the reporter cell line. HeLa-Cas9-poGFP/mCherry reporter cells were infected with the sgRNA targeting PEX3 for 48 hours, and subsequently treated with puromycin for two days. GFP/mCherry signals were analyzed by flow cytometry. The reporter cells infected with a non-targeting sgRNA (sgNTC) were used as a control. **(C)** Schematics of the screening procedure. HeLa-Cas9-poGFP/mCherry cells were transduced with the ubiquitin ligase lenti-CRISPR library. 24 hours after transduction, hygromycin were added to cells and maintained for 5 days. The surviving cells were sorted for the lowest and highest 30% of the GFP/mCherry ratio. sgRNAs were amplified from the extracted genomic DNA of each sample for deep sequencing. **(D)** Illustration of the top 10 candidate genes with highest Robust Rank Aggregation (RRA) scores, calculated from the enrichment of sgRNAs in the GFP low (GFP/mCherry ratio at the bottom 30%) cells compared to GFP high (GFP/mCherry ratio at the top 30%) cells (**Supplementary Table 3**). **(E)** Illustration of the top 10 candidate genes in the two-round sorting screen. RRA scores are calculated from the enrichment of sgRNAs in the sorted cells compared to unsorted cells.

To validate HeLa-Cas9-poGFP/mCherry as a reliable peroxisome function reporter cell, we designed a sgRNA to target *PEX3* (**Supplemental Table 1**). PEX3 is a peroxisomal protein required for the assembly of peroxisomal membrane proteins (PMPs) for the biogenesis of peroxisomes. *PEX3* deficient cells lack functional peroxisomes. Flow cytometry analysis of cells in the presence of sgRNA targeting PEX3 (sgPEX3) revealed a significant decrease in GFP/mCherry signal comparing to control cells (**Fig. 1B**). This suggests that HeLa-Cas9-poGFP/mCherry can be utilized as a reporter for peroxisome function.

We screened a specialized CRISPR sgRNA library, in which there are 5204 sgRNAs targeting 573 human genes (∼10 sgRNAs/each target) (**Supplemental Table 2)**. Each gene encodes a potential ubiquitin ligase, characterized with a RING domain, a HECT domain or a cullin-interacting domain. Amount those ubiquitin ligases, PEX2, PEX10 and PEX12 are three ubiquitin ligases that form a trimeric complex essential for peroxisome protein import.

We applied two types of screening strategies with different stringency, aiming to identify positive regulators. First, HeLa-Cas9-poGFP/mCherry cells were transfected with the sgRNA library, cultured with hygromycin selection for 5 days, then harvested and subjected for flow cytometry sorting. Cells with GFP/mCherry ratio at the bottom 30% and top 30% were sorted and sequenced individually (**Fig. 1C**). sgRNAs targeting positive regulators are expected to be enriched in the GFP low sample. Therefore, fold changes of sgRNAs (GFP low/GFP high) were analyzed (**Fig. S2**) and presented via Robust Rank Aggregation (RRA) score. Ubiquitin ligase PEX2, a well-known peroxisome transport regulator, ranks the highest in the screen (**Fig. 1D**). However, *PEX10* and *PEX12*, which are two ubiquitin ligases that form a trimeric complex with PEX2, did not appear as top-ranked hits in this one-round screen.

We also applied a more stringent two-round screening strategy. In this screen, cells with GFP fluorescence at bottom 30% were sorted and cultured for another 6 days, followed by another round of sorting cells with bottom 30% GFP intensity. The cells were then harvested for sgRNA sequencing. sgRNAs that target *PEX2, PEX10* and *PEX12* were all highly enriched in the sorted sample compare to unsorted cells (**Fig. 1E**). However, since PEX2, PEX10, and PEX12 are all essential for peroxisomal protein import, they dominated in the stringent screen. We rationalize that potential peroxisome regulators are more likely to be present in the less stringent one-round screening.

### ZBTB17 regulates peroxisome protein import

To validate the screening hits, we generated another reporter cell stably expressing Ub-GFP-SKL, in which ubiquitin is fused to the N terminus of GFP-SKL with Gly76 mutated to valine. Ub-GFP-SKL is fast degraded by the ubiquitin-proteasome degradation system in the cytosol (Dantuma et al., 2000), while it is protected in peroxisomes once imported. With this design, minimal cytosolic Ub-GFP-SKL can be detected, further eliminating non-peroxisomal GFP background.

Targeting top 10 hits from the screening, we transfected cells with individual sgRNAs targeting each gene and examined the fluorescence intensity of Ub-GFP-SKL. Among all validated genes, knocking out *ZBTB17* (also known as *MIZ-1*) significantly reduces the Ub-GFP-SKL intensity in cells, similar to what was observed for *PEX2, PEX10* and *PEX12* (**Fig. 2A**). The results were further confirmed in cells expressing GFP-SKL (**Fig. 2B**). The presence of sgRNA is indicated by the co-expressed mCherry. In the presence of a non-targeting sgRNA control (sgNTC), GFP-SKL mainly localizes within peroxisomes, showing GFP dots. In the presence of sgRNA targeting *PEX2*, *PEX10* or *PEX12*, the peroxisome protein import is impaired, resulting in the distribution of GFP-SKL in the cytosol. With the sgRNA targeting *ZBTB17*, the cells also display cytosolic GFP-SKL distribution, suggesting peroxisome defects (**Fig. 2B**).

**Figure 2.**
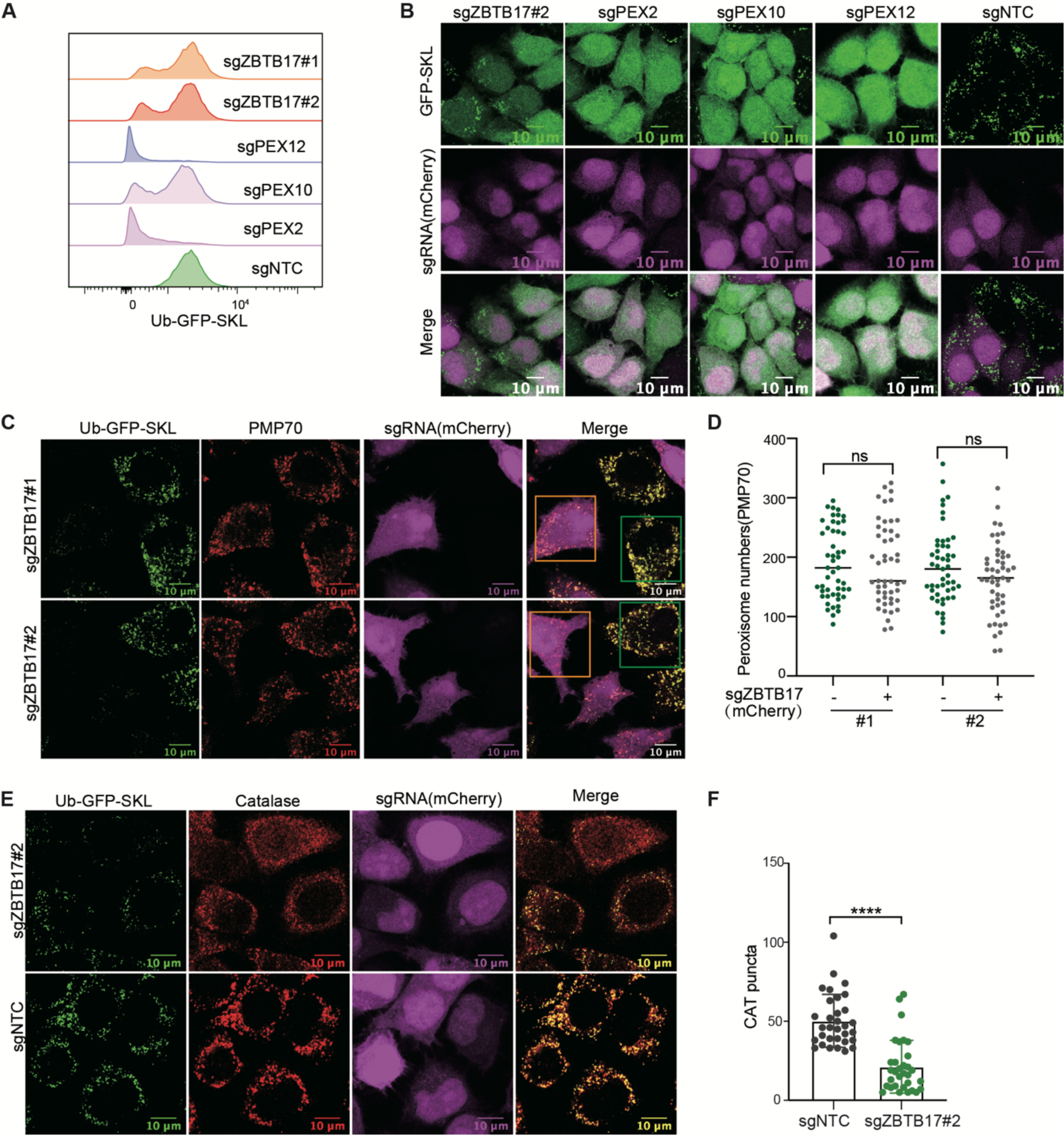
ZBTB17 regulates the translocation of peroxisomal matrix proteins. **(A)** ZBTB17 knockout reduces Ub-GFP-SKL levels. Flow cytometry analysis of GFP signals in HeLa cells stably expressing Ub-GFP-SKL (UGS cells in brief). sgZBTB17, sgPEX2, sgPEX10, sgPEX12 or sgNTC were used to knock out indicated genes. sgZBTB17#1 and sgZBTB17#2 are two different sgRNA sequences. Cells were analyzed 4 days after transfection. **(B)** ZBTB17 knockout causes the cytosolic retention of GFP-SKL. Fluorescence microscopy images showing the cellular location of GFP-SKL and mCherry in HeLa cells stably expressing GFP-SKL and transfected with different sgRNAs (sgZBTB17, sgPEX2, sgPEX10, sgPEX12). sgRNAs were co-expressed with mCherry. **(C)** Representative immunofluorescence microscopy images of Ub-GFP-SKL HeLa cells transfected with indicated sgRNAs and stained for PMP70. Orange box, sgRNA transfected cell, mCherry positive; Green box, un-transfected cell, mCherry negative. **(D)** Quantification of peroxisome numbers (PMP70 puncta) in > 90 cells for (C). **(E)** ZBTB17 knockout causes the cytosolic retention of catalase. Immunofluorescence microscopy images of Ub-GFP-SKL HeLa cells transfected with indicated sgRNAs and stained for catalase. Scale bars, 10 μm. **(F)** Quantification of Catalase puncta in (E). Values are means ± SD, calculated using 32 cells. ****p < 0.0001 by two-tailed Student’s t-test.

The abnormal GFP-SKL distribution can be explained by either defective GFP-SKL import or the loss of peroxisomes. Therefore, we examined peroxisome numbers. We transfected Ub-GFP-SKL containing cells with sgRNA targeting *ZBTB17* (mCherry positive) with low transfection efficiency, so the un-transfected cells (mCherry negative) can be used as the control group. The peroxisome membrane protein marker PMP70 was stained. The number of peroxisomes was determined by counting PMP70-positive specks, and statistical analysis revealed no change of peroxisome numbers in ZBTB17 knockout cells (**Fig. 2C and 2D**).

In addition to GFP-SKL, the cellular localization of catalase, an endogenous peroxisome matrix enzyme, was investigated. Catalase is widely distributed in the cytoplasm in the presence of sgRNA targeting *ZBTB17* (**Fig. 2E**). We also conducted a cellular fractionation experiment (**Fig. S3A**). In *ZBTB17* knockout cells, less catalase was detected in the peroxisome-enriched fraction (23k), accompanied by a decrease of PEX13 level in these fractions (**Fig. S3B**). Catalase activity was measured, suggesting a reduced enzyme content in peroxisomes and an increase in the cytosol in *ZBTB17* knockout cells (**Fig. S3D**). PMP70 levels remained consistent in different cells (**Fig. S3B**), aligning with the observation of unchanged peroxisome numbers. Therefore, ZBTB17 regulates peroxisome protein import.

### The ubiquitin ligase activity of ZBTB17 is not required for peroxisome protein import

ZBTB17 contains a BTB domain at the N terminus and 13 zinc fingers (ZNFs) at the C terminus (**Fig. 3A**). It is included in the screening library as a ubiquitin ligase because the BTB domains, in general, bind Cul3 to form BTB/Cul3 ubiquitin ligases. ZBTB17 also functions as a transcription factor, but mainly via the C terminal ZNFs. We first sought to determine whether ZBTB17 plays a role as a ubiquitin ligase in peroxisome protein import.

**Figure 3.**
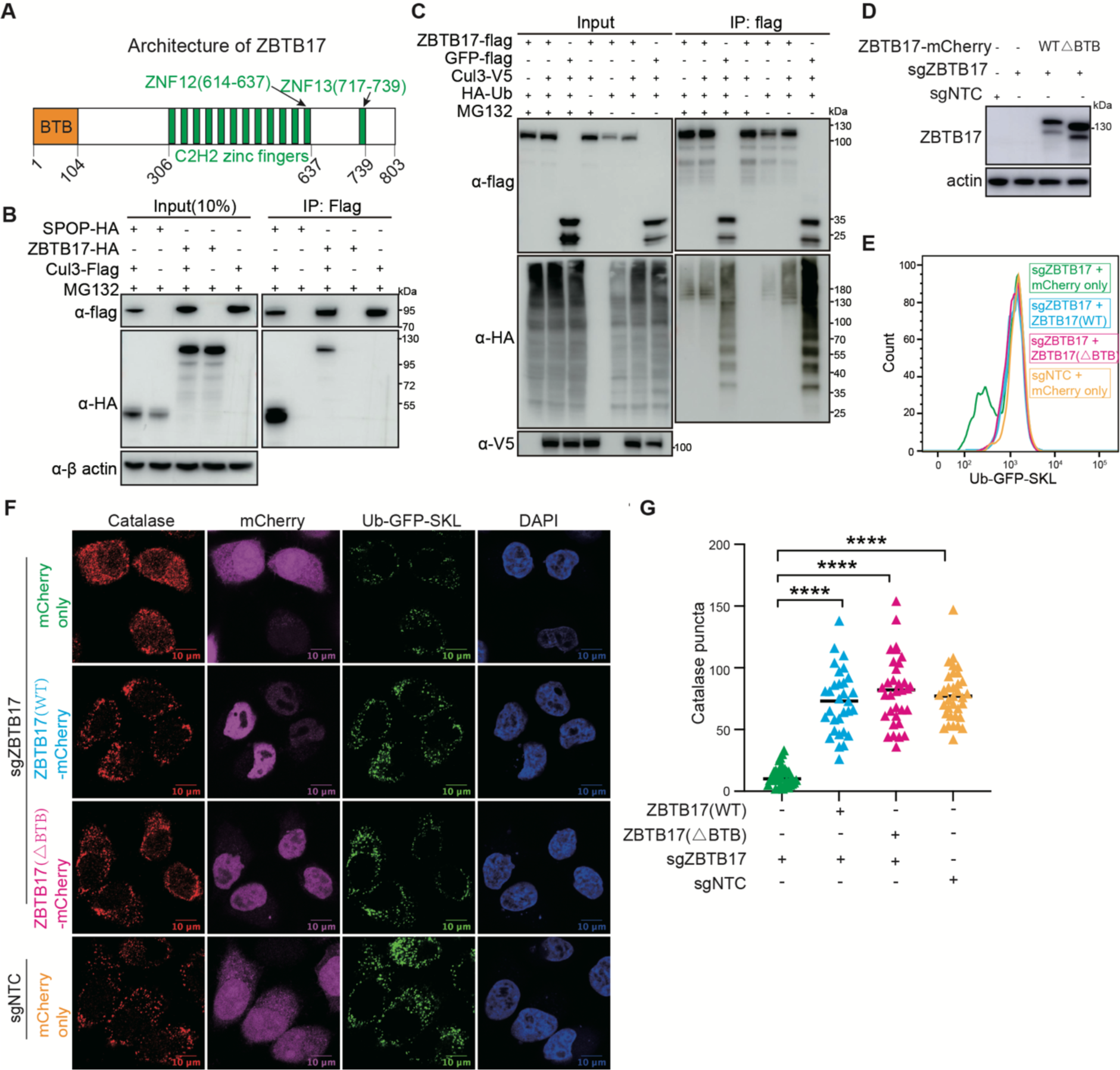
The BTB domain of ZBTB17 is not required for peroxisome protein import. **(A)** Schematics of the domain arrangement of ZBTB17. **(B)** Co-immunoprecipitation of ZBTB17-HA or SPOP-HA with Cul3-Flag in HEK293T cells. HEK293T cells were co-transfected with either ZBTB17-HA or SPOP-HA along with Cul3-Flag. Cell lysates were then subjected to immunoprecipitation using an anti-Flag antibody, followed by immunoblotting with the indicated antibodies. **(C)** Ubiquitination assay of ZBTB17 in cells co-transfected with HA-Ub, ZBTB17-Flag, GFP-Flag, and Cul3-V5. HEK293T cells were transfected with the mentioned plasmids for 48 h and subsequently treated with 10 μM MG132 for 6 h. Cell lysates were immunoprecipitated using an anti-Flag antibody and then analyzed by immunoblotting with the respective antibodies. **(D)** Immunoblotting analysis of ZBTB17 protein expression. **(E)** Analysis of Ub-GFP-SKL signal in ZBTB17 knockout UGS cells. ZBTB17 was eliminated from UGS cells using sgRNA. Following infection with ZBTB17 (WT)-mCherry, ZBTB17 (△BTB)-mCherry, or mCherry-only virus for 48 hours, the Ub-GFP-SKL signal was assessed via flow cytometry. **(F)** Immunofluorescence (IF) analysis of catalase and Ub-GFP-SKL. ZBTB17 was eliminated from UGS cells using sgRNA. Following this, cells were infected with ZBTB17 (WT)-mCherry, ZBTB17 (△BTB)-mCherry, or mCherry-only virus for 48 hours. The scale bar represents 10 μm. **(G)** Quantification of catalase puncta in UGS cells. Calculations were made using data from (E) (> 30 cells for each condition).

To address if ZBTB17 forms an active ubiquitin ligase with Cul3, we examined the interaction between ZBTB17 and Cul3. Exogenously expressed ZBTB17 with HA tag associates with Flag tagged Cul3 (**Fig. 3B**). However, the interactions between ZBTB17 and Cul3 seems to be much weaker than the interaction between the classic Cul3-binding BTB protein SPOP and Cul3, evidenced by less ZBTB17-HA are immunoprecipitated with Cul3-Flag. However, the weak interaction between ZBTB17 and Cul3 is sufficient to mediate the self-ubiquitination of ZBTB17 in a Cul3 dependent manner (**Fig. 3C**).

To address if the BTB domain is required for normal Ub-GFP-SKL signals in cells, we re-introduced wildtype ZBTB17 or ZBTB17(τιBTB) into *ZBTB17* knockout cells (**Fig 3D**). Full-length ZBTB17 reconstitutes the GFP signal as expected. The expression of ZBTB17(τιBTB) is also sufficient to recover the GFP signal (**Fig. 3E**), suggesting the BTB domain of ZBTB17 is not a major contributor for peroxisome regulation. The import of Ub-GFP-SKL and catalase into peroxisomes were recovered when wildtype ZBTB17 or ZBTB17(τιBTB) was expressed in *ZBTB17* knockout cells (**Fig. 3F** and **3G**). These data suggest although ZBTB17 may have function as a ubiquitin ligase, the ubiquitin ligase activity of ZBTB17 is not required to maintain normal peroxisome import.

### ZBTB17 regulates *PEX13* transcription

In addition to the role as a ubiquitin ligase, ZBTB17 has been characterized for its role as a transcriptional regulator (Kosan et al., 2010; Peukert et al., 1997b; Wiese et al., 2013). ZBTB17 mainly localizes in the nuclei (**Fig. 3F**). Therefore, we investigated ZBTB17 as a transcriptional regulator for peroxisome protein import.

HeLa cells were stably transfected with either a non-targeting shRNA or an shRNA specifically targeting ZBTB17. Transcriptome analysis based on RNA sequencing, was carried out in HeLa cells with or without ZBTB17 knockdown. A comparative analysis of gene expression yields a total of 3178 differentially expressed genes (**Supplementary Table 4**). Remarkably, among these genes, 15 genes associated with the peroxisome pathway were identified and their roles in peroxisome function are highlighted in the KEGG (Kyoto Encyclopedia of Genes and Genomes) pathway (**Fig. S4**). The heatmap displays a significant alteration in the expression of multiple peroxisome-related genes in cells with *ZBTB17* knockdown (**Fig. 4A**). Specifically, five genes exhibited upregulated expression, while ten genes displayed downregulated expression. Notably, PEX13, one of the key peroxins that control the peroxisomal protein import, has been down-regulated. The relative ZBTB17 and PEX13 transcripts in cells were derived from the RNA-seq data, confirming the down-regulation of PEX13 (**Fig. S5A**).

**Figure 4.**
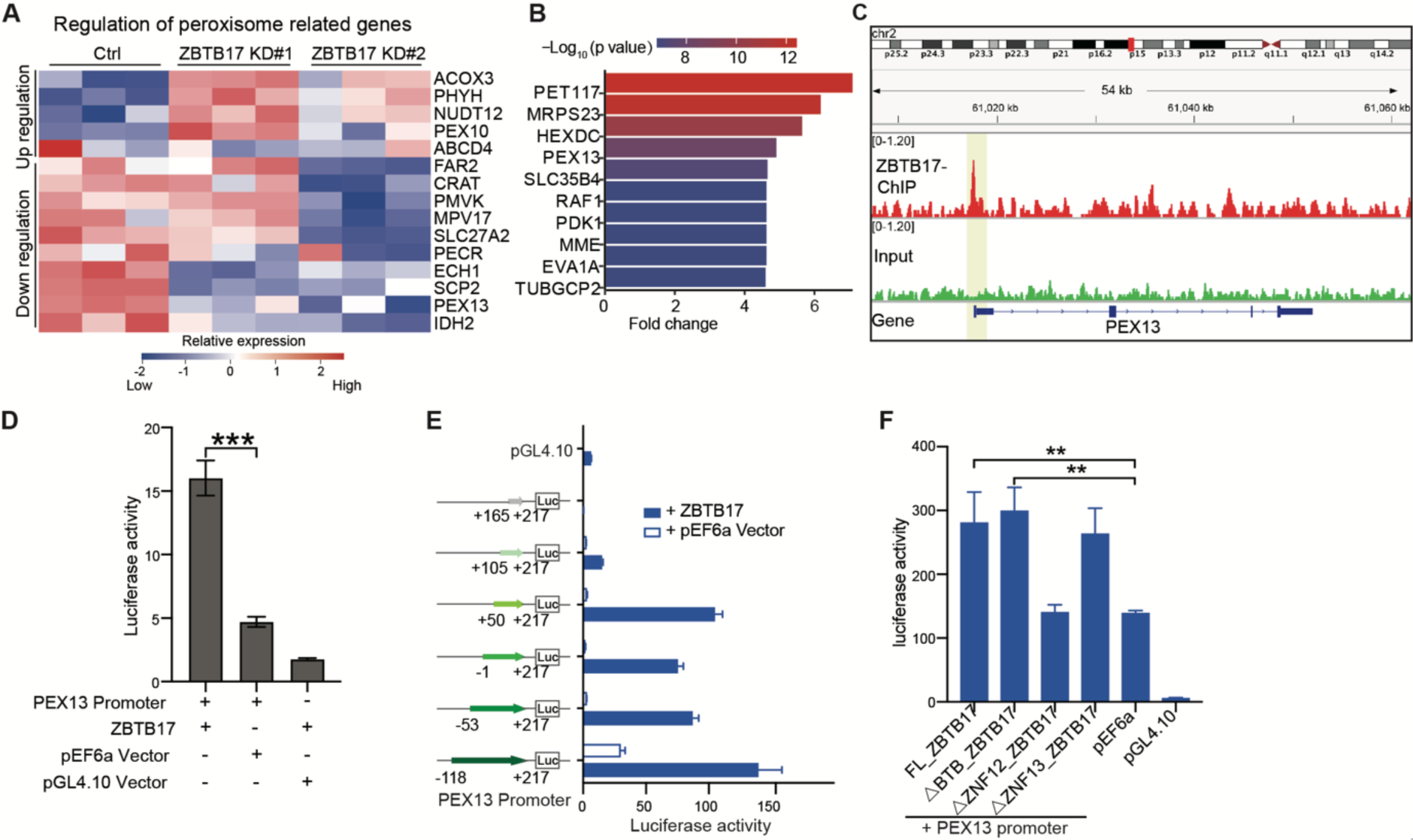
ZBTB17 regulates PEX13 transcription directly by associating with PEX13 promoter region. **(A)** Heat map of differentially expressed peroxisomal genes in response to ZBTB17 knockdown. HeLa cells were infected with ZBTB17 shRNA (shZBTB17) or non-targeting shRNA (shNT) for 4 days, and were subjected to RNA-seq analysis. Lower and higher levels of expression are presented in shades of blue and red, respectively (fold change > 1.3; p.adjust < 0.05). **(B)** Analysis of correlation between genes bound by ZBTB17 in ChIP-seq data and genes that are down-regulated in RNA-seq data. The top ten genes are displayed, sorted by p-value. **(C)** ChIP-seq data from HeLa cells showing ZBTB17 occupancy at the PEX13 promoter. **(D)** Luciferase assay demonstrating the interaction of ZBTB17 with the PEX13 promoter. HEK293T cells were co-transfected with pGL4.10-PEX13 promoter (−1886 to +115)-dual luciferase reporter and pEF6a-PEX13 for 24 hours. Luminescence was quantified using a Luminescence Counter. Empty pGL4.10 and pEF6 vectors were used for controls. **(E)** Luciferase assay to map ZBTB17 binding regions on PEX13. The PEX13 promoter sequence (335 bp) was truncated into shorter fragments and cloned into the luciferase reporter vector. Luminescence was quantified in the presence or absence of con-transfected ZBTB17. **(F)** Luciferase assay to map PEX13 promoter binding regions on ZBTB17. △BTB, △ZNF12, △ZNF13 are ZBTB17 with the BTB (residues 1-104), ZNF12 (residues 614-637), or ZNF13 (residues 717-739) removed individually.

PEX13 interacts with other peroxins, such as PEX5 and PEX14, forming a dynamic protein complex that aids in the translocation of proteins across the peroxisomal membrane (Barros-Barbosa et al., 2019; Gao et al., 2022; Meinecke et al., 2010; Ravindran et al., 2023). In addition to PEX13, peroxisomal matrix protein translocation also relies on PEX2, PEX5, PEX7, PEX10, PEX12, PEX14, and PEX26. Therefore, we examined the transcriptional level of these import-related genes individually by real time RT-PCR (reverse transcription– polymerase chain reaction) in *ZBTB17* knockdown cells (**Fig. S5B**). Most gene transcripts are not affected by *ZBTB17* knockdown, while the mRNA level of *PEX13* is significantly downregulated. *PEX2* expression is slightly reduced, but not to the extent as *PEX13*.

To further confirm the role of ZBTB17 as a transcriptional regulator, we mapped the associated DNA sequence by chromatin-immunoprecipitation (Ch-IP). By combining ChIP-seq and RNA-seq data, a few key genes are identified as potential ZBTB17 direct targets (**Fig. 4B**). *PEX13* ranks high, and the ChIP-seq data also revealed that ZBTB17 binds specifically to the promotor region (nt -118 to +217) of *PEX13* (**Fig. 4C**). However, other peroxisomal genes, such as *PEX2*, were not identified, suggesting that *PEX13* is the primary target gene and the others are likely being regulated indirectly.

To further confirm *PEX13* as the direct target of ZBTB17, we performed the luciferase assay. In 293T cells, the insertion of the *PEX13* promoter in a luciferase expression vector induced luciferase expression, the presence of exogenously expressed ZBTB17 further enhanced luciferase activity (**Fig. 4D**). To narrow down the ZBTB17 binding region on *PEX13* promoter, we performed a systematic mapping. With the 335 bp sequence covered by ChIP-seq, the presence of ZBTB17 significantly induced luciferase expression. Systematic truncations identified NT +50 to +105 on *PEX13* promoter region are critical for ZBTB17 trans-activity (**Fig. 4E**). ZBTB17 contains 13 ZNFs and it was previously reported that ZNF12 and ZNF13 are important for DNA binding (Boisvert et al., 2022; Peukert et al., 1997a; Tremblay et al., 2016). In case of *PEX13* transcriptional regulation, we delete ZNF12 and ZNF13 individually. The *ΔZNF12* mutant is defective to induce luciferase activity, suggesting ZBTB17 binds *PEX13* promoter via the ZNF12 motif (**Fig. 4F**).

Some BTB domains also bind DNA, as suggested in other BTB domain-containing proteins, such as BCL6 and PLZF (Stogios et al., 2005). However, the truncation of *ZBTB17* BTB domain had no effect on *PEX13* transcription (**Fig. 4F**). This result is also consistent with the observation that ZBTB17(ΔBTB) can rescue peroxisome protein import in *ZBTB17* KO cells (**Fig. 3E**).

### ZBTB17 regulates peroxisome protein import mainly *via PEX13*

Consistent with the mRNA level, the protein level of PEX13 is also significantly decreased in cells with *ZBTB17* KD (**Fig. 5A**). As PEX13 is one of the key players for peroxisomal protein import, we sought to investigate whether the restoration of PEX13 protein level alone is adequate to re-establish peroxisomal protein import. In Ub-GFP-SKL cells, *ZBTB17* were knocked down with shRNA, mCherry fused PEX13 or mCherry alone were introduced into these cells, and the expression of ZBTB17 and PEX13 in each sample were confirmed by western blots (**Fig. 5B**). The Ub-GFP signal was quantified in each sample as an indicator of peroxisomal import. GFP signal was decreased in *ZBTB17* knockdown cells, but was restored upon exogenous expression of PEX13-mCherry, while mCherry alone had no effect on the GFP signal (**Fig. 5C**). Similarly, we examined the localization of Ub-GFP-SKL and catalase under these conditions. The exogenous expression of PEX13 alone is adequate to restore peroxisomal import in the absence of ZBTB17 (**Fig. 5D** and **5E**). Despite the downregulated mRNA expression of *PEX2* (**Fig. S5B**), these results again suggest that *PEX13* is the primary target of ZBTB17 in regulating peroxisomal import.

**Figure 5.**
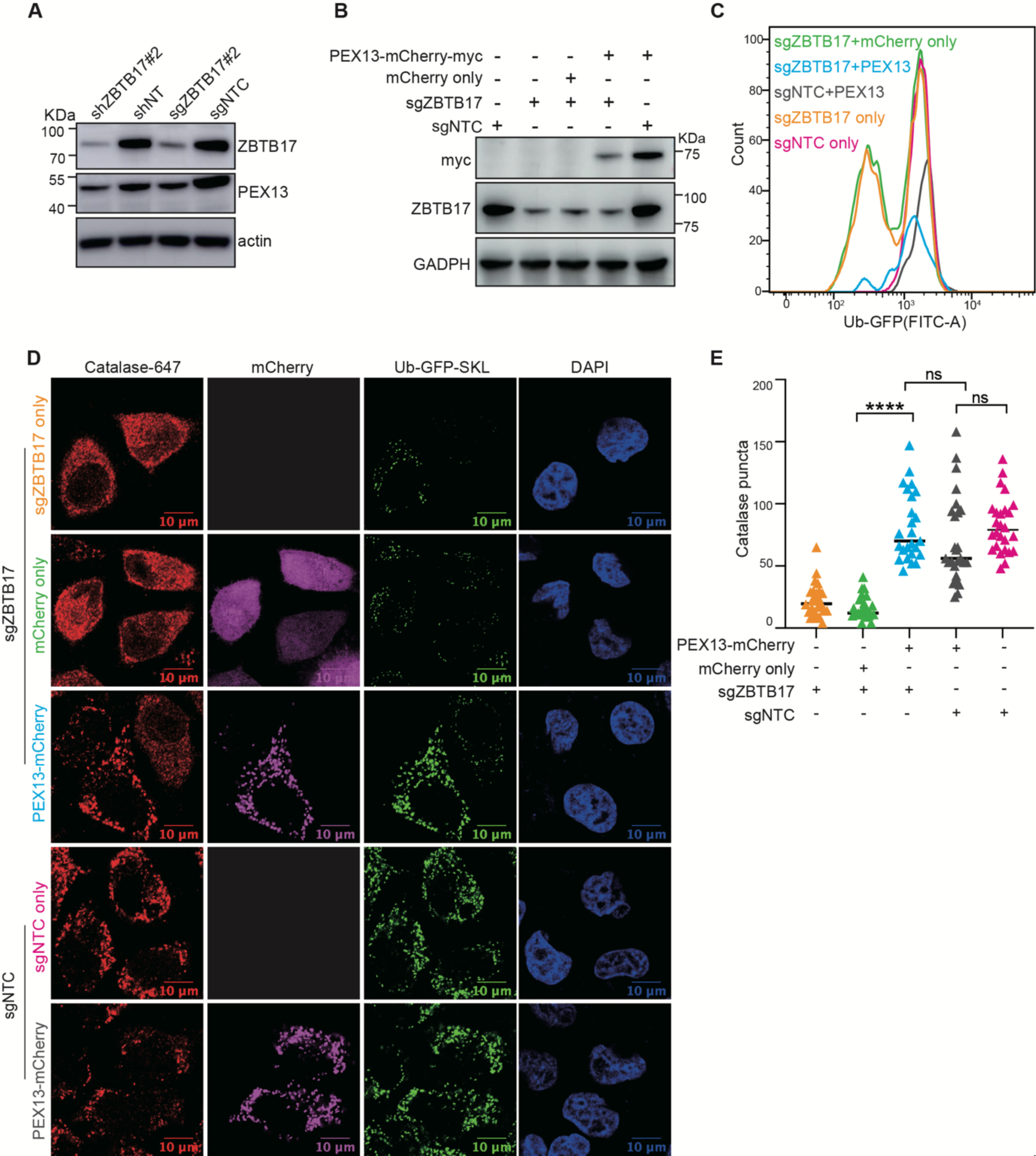
ZBTB17 regulates peroxisome protein import via PEX13. **(A)** Immunoblotting (IB) of PEX13 and ZBTB17 in HeLa cells following with shRNA or sgRNA/Cas9 targeting ZBTB17. **(B)** Immunoblotting (IB) showing the corresponding protein expression levels in (C). **(C)** Flow cytometry analysis of Ub-GFP-SKL in ZBTB17 knockout HeLa cells expressing either PEX13-mCherry-myc or mCherry for 48h. **(D)** Representative immunofluorescence images showing the localization of catalase in corresponding conditions in (B). Scale bars represent 10 μm. **(E)** Quantification of number of catalase puncta in (D). Calculated using > 25 cells/sample.

### ZBTB17 modulates purine metabolism through its transcriptional regulation of peroxisome import

Peroxisomes are involved in the catabolism of lipids, D-amino acids, polyamines, purines, as well as reactive oxygen species (ROS) metabolism. Dysregulation in the import of peroxisome matrix proteins may impact these peroxisome-dependent metabolic pathways.

Since PEX13 is regulated by ZBTB17, we wonder if ZBTB17 affects peroxisome-dependent metabolic pathways. We profiled the metabolomic changes in *ZBTB17* or *PEX13* knockdown cells using liquid chromatography-mass spectrometry (LC-MS)-based untargeted metabolomics. Metabolite annotation was performed using MetDNA (Shen et al., 2019b; Zhou et al., 2022b) and metabolomics data was normalized to sample protein concentrations. Metabolite peak intensity was used to calculated the fold changes to indicate the metabolic changes. Comparing to the control cells, 8 metabolites in *ZBTB17* knockdown cells and 10 metabolites in *PEX13* knock down cells were significantly down-regulated (p<0.05; and fold change>2) (**Fig. 6A and B, Supplementary Table 5**). Metabolic pathway enrichment analysis further revealed that the purine metabolism pathway was one of the down-regulated pathways in both *ZBTB17* and *PEX13* knockdown cells (**Fig. 6C** and **6D**). Ribothymidine, inosine, inosine 5’-monophosphate (IMP) and guanosine were remarkably reduced in both *ZBTB17* and *PEX13* knockdown cells. Hypoxanthine was only decreased in *ZBTB17* knockdown cells, while Guanosine 5’-monophosphate (GMP) were decreased in *PEX13* knockdown cells (**Fig. 6E** and **6F**). These data show the consistent metabolic changes in *ZBTB17* and *PEX13* knockdown cells, suggesting ZBTB17-mediated metabolic changes are mainly contributed by the down-regulation of PEX13 and the consequent defect of peroxisome function.

**Figure 6.**
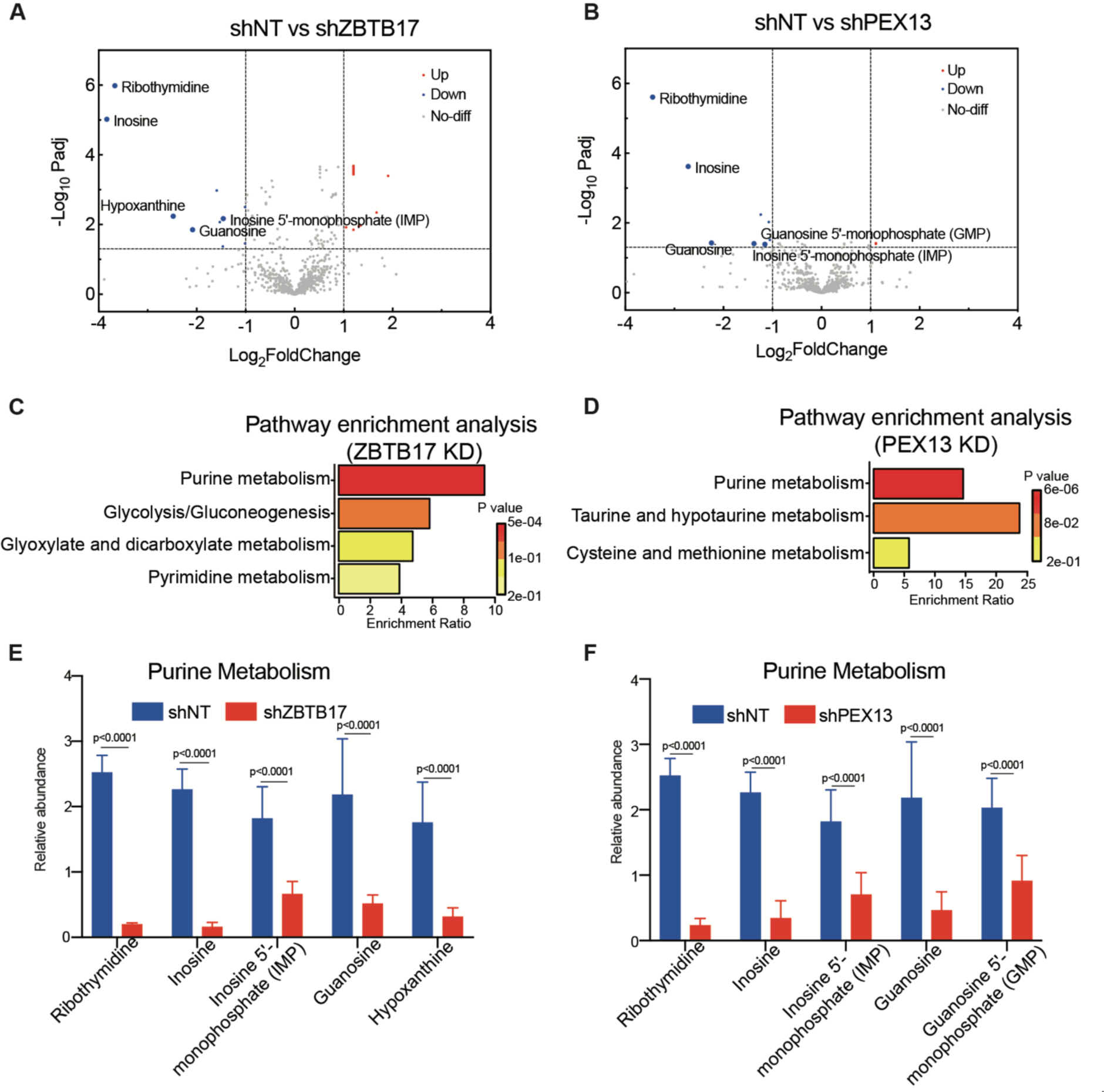
Metabolic alterations in cells with ZBTB17 or PEX13 knockdown. **(A)** The volcano plot of the fold changes of metabolites in cells with or without ZBTB17 knockdown. P values were determined by a two-tailed Student’s t-test. Blue dots indicate significantly changed metabolites (p < 0.05 and fold change >2, n = 6 biologically independent samples in each group). **(B)** The volcano plot of the fold changes of metabolites in cells with or without PEX13 knockdown. P values were determined by a two-tailed Student’s t-test. Blue dots indicate significantly changed metabolites (p < 0.05 and fold change >2, n = 6 biologically independent samples in each group). **(C)** and **(D)**, Significantly decreased metabolites in ZBTB17 or PEX13 knockdown cells are enriched in the purine metabolism pathway. MetaboAnalyst (https://www.metaboanalyst.ca) were used for pathway enrichment analysis. **(E)** and **(F)**, Relative abundances of each purine metabolism-related metabolite in cells with ZBTB17 or PEX13 knockdown are illustrated. Values are mean ± SD, n = 6 independent samples in each group. P values were determined by two-way ANOVA multiple comparisons test.

## DISCUSSION

We have presented here a focused genetic screen employing the CRISPR/Cas9 system to identify novel regulators of peroxisomes in cells. Our screen has generated a comprehensive dataset that not only reaffirms the role of known peroxisomal genes, but also unveils previously unrecognized transcriptional regulation mechanism of peroxisomes, and the related purine metabolism.

Despite the power of the CRISPR-based screening system, our screens suggest the selection strategy matters for peroxisomes. Cell viability and the fluorescence reporter are two major phenotypic readouts. A recent study used cell viability to screen peroxisome regulators, and revealed the connection between Wnt signaling and peroxisomes (Vu et al., 2024). In this study, we used fluorescence reporter, which can be fine-tuned to adapt different screening stringency. In this study, we arbitrarily set bottom 30% GFP/mCherry expression as the cut-off. One or two rounds of selection were used. For future screening of more comprehensive sgRNA libraries, different selection strategies with different fluorescence cutoff or selection rounds should be tested and applied to best fit the needs. In general, a stringent screen is more likely to identify the dominant regulators, while a less stringent screen will enhance the possibility to identify all regulators, but also inevitably increase the chance for false positive enrichment.

While we have identified ZBTB17 as one of the regulators of peroxisomal protein import, it’s important to note that the GFP-SKL/mCherry reporter system is versatile. In fact, changes in the GFP fluorescent signal may reflect alterations in peroxisome synthesis, peroxisome protein import, or peroxisome degradation. Therefore, when utilizing the GFP-SKL/mCherry reporter as an indicator of peroxisome function, additional assays are often necessary to comprehensively assess which aspects of peroxisome function have been affected.

Our screen identified ZBTB17, a previously unrecognized regulator of the peroxisomal protein PEX13, as a positive regulator of peroxisome import. ZBTB17 belongs to the BTB ubiquitin ligase family, characterized by its N-terminal BTB domain. We confirmed the interaction between ZBTB17 and Cullin3, a scaffold protein for cullin-RING ligases (CRLs), and assessed the potential ubiquitin ligase activity of ZBTB17. However, we discovered that the BTB domain is dispensable for ZBTB17-mediated transcriptional regulation of PEX13. The role of ZBTB17 as a transcriptional regulator may have been underestimated, as previous *ZBTB17* knockout mice were created by deleting the BTB/POZ domain (Do-Umehara et al., 2020). Our discovery that ZBTB17 without the BTB domain retains some of its function as a transcription factor implies that different *ZBTB17* knockout strategies should be considered.

PEX13 appears to be a pivotal node in the transcriptional regulation of peroxisome import. Besides PEX13, the expression of several other genes related to peroxisome import was also affected. Notably, PEX2 was down-regulated, while PEX14 exhibited up-regulation (**Fig. S5B**). Despite the upregulation of PEX14, the absence of PEX13 resulted in a restriction of protein import, in line with the indispensable role of PEX13 in protein translocation (Elgersma et al., 1996; Erdmann and Blobel, 1996; Gould et al., 1996). Apart from the genes involved in the peroxisomal import machinery, several peroxisomal proteins, including metabolic enzymes like SCP2, ECH1, and PECR exhibit downregulation (**Fig. 4A**). This observation suggests that ZBTB17 may have a broader impact on diverse metabolic pathways within peroxisomes, a dimension not investigated in our current study. It is conceivable that ZBTB17 governs peroxisome function by influencing both protein import and the expression levels of metabolic enzymes.

In the absence of ZBTB17, the transcription of PEX13 is not fully blocked but decreased (**Fig. 5A**). A recent study shows that pexophagy is induced in the absence of PEX13 (Demers et al., 2023). We have monitored peroxisome numbers and PEX13 protein levels over time (**Fig. S6**). Pexophagy is usually triggered by the deficiency of peroxisome import. However, peroxisome numbers in *ZBTB17* knockout cells do not change over 10 days, despite the import defect (**Fig. S6A, S6B**). In *ZBTB17* knockout cells, the down-regulation of PEX13 is accompanied by the down-regulation of peroxisomal enzymes, suggesting less importing stress in ZBTB17 deficient cells than in PEX13 deficient cells. In addition, PEX13 protein level in ZBTB17 deficient cells is not fully ablated over time (**Fig. S6C, S6D**). We speculate there are other factors involved in PEX13 transcriptional regulation. In fact, it has been reported that ZBTB17 partners with many other transcriptional factors to either enhance or inhibit gene transcription (Aesoy et al., 2014; Wiese et al., 2013). ZBTB17 is one transcriptional regulator that fine-tunes the expression of PEX13, thereby impacting peroxisome import and influencing cellular metabolism.

Peroxisomes are closely associated with purine metabolism, as they house three essential enzymes—xanthine dehydrogenase (XDH), Uricase (Uox), and allantoicase (ALLC)— involved in purine degradation, all located within the peroxisome matrix (Wanders, 2014). However, in human cells, active Uox and ALLC are absent, leading to an inability to further oxidize urate. XDH serves as a crucial peroxisomal enzyme in purine metabolism, catalyzing the oxidation of hypoxanthine to xanthion and xanthion to urate. Any defect in peroxisomal import can result in the mislocalization of XDH, impacting the concentration of hypoxanthine and other upstream metabolites in the same pathway, including inosine, inosine 5’-monophosphate (IMP), and guanosine. As a result, it is not surprising that purine metabolism is downregulated in PEX13 knockdown cells. Our study also demonstrates a similar downregulation of purine metabolism in ZBTB17 knockdown cells, further corroborating that the metabolomic changes induced by the transcription factor ZBTB17 are related to its regulation of peroxisomal import via PEX13. Peroxisomal homeostasis can be regulated through transcription factors such as PPAR family proteins and HIF transcription factors (Lemberger et al., 1996; Walter et al., 2014). We have revealed one more transcriptional regulation mechanism for peroxisome function.

## METHODS

### Cells and reagents

HeLa and HEK293 FT cells were cultured in Dulbecco’s modified Eagle’s medium (Gibco, C11995500BT-20) supplemented with 10% fetal bovine serum (Gemini,100-500),1% penicillin-streptomycin (Gibco, 15140122) and 5% CO_2_ in 37 °C.

Antibodies and other reagents used: anti-Flag (GNI4110-FG-S 1:1000 WB, GNI), anti-Flag-HRP (GNI4310-FG-S 1:1000 WB, GNI), anti-HA (2367S 1:1000 WB, Cell Signaling), anti-HA-HRP (2999S 1:1000 WB, Cell Signaling), anti-V5 (13202S 1:1000 WB, Cell Signaling), anti-actin (60008-1-Ig-100ul 1:1000 WB, Proteintech), anti-GAPDH (AP0063-50ul 1:1000 WB, Bioworld), anti-Tubulin (AC012 50 μL 1:1000 WB, ABclonal), anti-PMP70 (P0497-200UL 1:500 IF, Sigma), anti-PMP70 (SAB4200181-200UL 1:1000 WB, Sigma), anti-Catalase (219010-1ML 1:1000 WB, 1:500 IF, Merck), anti-PEX13 (sc-271477 1:500 WB, Santa Cruz), anti-ZBTB17 (sc-136985 1:500 WB, Santa Cruz), anti-ZBTB17 (sc-136985X ChIP, Santa Cruz), Donkey Anti-Rabbit IgG H&L (Alexa Fluor® 647) (ab150075 1:500 IF, Abcam), Hygromycin (ant-hg-1,Invitrogen), puromycin (ant-pr-1,InvivoGen), polybrene (40804ES86,Yeasen).

### Generation of the peroxisome reporter GFP-SKL-IRES-mCherry cell line

To generate the peroxisome reporter cell line, we first generated the HeLa cell line stably expressing spCas9. Lentivirus was produced by co-transfecting Cas9-T2A-BFP-Flag plasmid along with two viral packaging plasmids into HEK293 FT cells using Polyethylenimine transfection reagent (PolySciences, 9002-98-6). Approximately 48 hours after transfection, the viral supernatant was harvested and subsequently filtered through a 0.22 μm filter. Additionally, to increase viral titer, the supernatant was concentrated using PEG virus precipitation reagents (BioVision, K904-50). HeLa cells were transfected with Cas9-T2A-BFP-Flag virus. Two days post-transduction, BFP-positive cells were sorted into a 96 well-plate for single cell clones using a BD FACS AriaIII flow cytometer. The single-cell clones were allowed to amplify for 2∼3 weeks. Immunoblot assay with the anti-Flag antibody was conducted to assess the expression of Cas9 and T7 endonuclease I-cutting assay (NEB, M0302S) was used to examine the activity of spCas9 in the isolated clones. Clone # 3-5-5 was used for future screening experiments. To generate the peroxisome reporter GFP-SKL-IRES-mCherry cell line, spCas9 expressing HeLa cells were transduced with a GFP-SKL-IRES-mCherry lentivirus. Cells expressing both GFP and mCherry were isolated by fluorescence-activated cell sorting using a BD FACS AriaIII. Single cell clones were generated and validated.

### CRISPR library Screen

A total of 1.8 x 10^7^ GFP-SKL-IRES-mCherry HeLa cells were transduced on Day 1 with the ubiquitin ligase lenti-CRISPR library (a gift from Haopeng Wang’s lab) at an MOI of 0.3 with an approximately 1000-fold library coverage. 24 hours after transduction, hygromycin (600 μg/ml) were added to cells and maintained for 5 days. On Day 7, the surviving cells were sorted for the lowest and highest 30% of the GFP/mCherry ratio by BD FACS Arial II.

To extract genomic DNA, cells were resuspended in 400 μl P1 buffer (Qiagen, 19051) with 100 μg/mL RNaseA and 40ul 10% SDS (MDBio, S001-100g). After incubate at room temperature for 15 min, the lysate was heated at 55°C for 30 min in the presence of Proteinase K (100 μg/mL, Sigma, P6556-25MG). After digestion, samples were passed through a needle of different sizes (21G-23G-25G-27G) for multiple times to shear DNA to ∼20kb size. (The criterion for judgment is whether the solution can easily pass through the needle). 400ul Phenol: Chloroform: Isoamyl Alcohol (25:24:1, v/v) was added into homogenized samples before the centrifugation at 3,000g for 20 min at room temperature. The aqueous phase was transferred into ultracentrifuge tubes and thoroughly mixed with 40 μl 3M sodium acetate plus 320 μl isopropanol at room temperature before centrifugation at 12,000g for 15 min at 4°C. The gDNA pellets were carefully washed with 1 mL 70% ethanol and centrifuged at 3,000g for 15 min. The pellets were dried at 37°C for 30 min and resuspended in water.

Multiple PCR reactions were prepared to amplify the region coding sgRNA from the extracted genomic DNA. For the first round of PCR, the total volume was 100 μL containing 50 μg sheared gDNA, 0.3 μM forward (5′-CTGCCATTTGTCTCGAGGTCG -3′) and reverse (5′-GCTCGGCGCCAGTTTGAATAT -3′) primer, 200 μM each dNTP, 1× Titanium Taq buffer and 1 μL Titanium Taq (Clontech, 639209). PCR cycles were: 1× (94°C 3min), 16× (94°C 30s, 65°C 10s, 72°C 20s), 1× (68°C 2min). A second round PCR were performed to add different barcodes for different samples. The total volume of the second round PCR was 100μL containing 2 μL round 1 PCR product, 0.5 μM forward and 0.5 μM reverse primer, 200 μM dNTP, 5× Prime Star buffer, and 1 μL Prime Star DNA Polymerase (Takara, R010A). PCR cycles were: 1× (94°C 3min), 16× (94°C 30s, 55°C 10s, 72°C 20s), 1× (68°C 2min). The main PCR products (∼250bp) were gel-purified from a 1% agarose gel and submitted for sequencing on an Illumina HiSEq-PE150. Sequencing reads were aligned to the sgRNA library and quantified by MaGeCK-count analysis 1. A pseudo count was added to each value and the log_2_-transformed fold change in abundance was then calculated between low and high fractions by MaGeCK-test analysis.(Li et al., 2014)

Primers used to add barcodes:

Forward, 5-AATGATACGGCGACCACCGAGATCTACACCGACTCGGTGCCACTTTT-3’;

Reverse, 5-CAAGCAGAAGACGGCATACGAGATCCTGAGATTTCTTGGGTAGTTTGCAGTTTT-3’ (GFP low population) and 5-CAAGCAGAAGACGGCATACGAGATCGACTCATTTCTTGGGTAGTTTGCAGTTTT-3’ (GFP high population)

### Gene knockout and knockdown

To knockout a specific gene in cells with CRISPR/Cas9, sgRNA targeting the specific gene was cloned into MP-783 (Tromp et al., 2018). Lentiviral vector expressing PuroR-T2A-mCherry under the EF1A promoter and a sgRNA sequence under the U6 promoter. MP-783 sgRNAs lentivirus production and infection were performed as described in above section. 2 days after infection, HeLa cells were treated with puromycin (2μg/mL) for 3 days for selection. The surviving cells were sorted for mCherry-positive by BD FACSAriaI II. The sgRNA sequences used in this study are summarized in the **Supplementary Table 1**.

To knockdown a specific gene, we used shRNA. The shRANs targeting a specific gene was cloned into pLKO.1 puro (10878; Addgene) vector using EcoRI (R3101, NEB) and AgeI (R3552S, NEB) restriction sites. HeLa cells were transfected with shRNA lentivirus (shZBTB17, shPEX13 and shNT) for 2 days, then add puromycin(2ug/mL) to select for infected cells. Change to fresh puromycin-containing media for 4 days. The sequences of shRNA used in this study are summarized in **Supplementary Table 6**.

### Immunofluorescence

HeLa cells were cultured overnight on glass coverslips in DMEM, supplemented with 5% CO2, at 37°C. The cells were rinsed three times with 1× phosphate-buffered saline (PBS) and fixed for 15 minutes at room temperature in 4% paraformaldehyde (meilunbio, MA0192). Cells were permeabilized with 0.1% NP-40 in PBS at room temperature for 15 min. Coverslips were then blocked with 2.5% BSA (Yeasen, 36105ES25) in Cell stanning buffer (4A Biotech, FXP005) for 1 hour at room temperature before adding primary antibody stain to coverslips at 4°C overnight. After primary staining, cells were wash 3 times with PBS and then stained with fluorochrome-conjugated secondary antibodies for 1.5-2 hour at room temperature. After stanning, Coverslips were incubated with DAPI for 10 min and then were adhered to a microscopy slide using Mounting Medium (Vectorlabs, H-1000). Coverslips were imaged on a Zeiss LSM 980 Airyscan2 (objective ×40 1.30 NA OIL).

### Immunoblot and immunoprecipitation

Cells were homogenized in lysis buffer (50 mM Tris, 200 mM NaCl, 1% NP-40, pH 7.5) with protease inhibitors (Biomake, B14001) for 30 min at 4°C. The lysates were centrifuged at 13,000g for 10 min at 4 °C to remove debris and to collect the whole cell extract. The supernatant was subjected to BCA Protein Assay (TIANGEN, PA115-01) to quantify protein levels. After boiling the sample with 1x Protein Loading Buffer (TransGen Biotech, DL101-02) at 95 °C for 10 min, equal amounts of proteins were loaded and separated on a 4-20% SDS–PAGE (GenScript, M42015C). The proteins were transferred to PVDF membrane (Cytiva, 10600023) and incubated with specific primary antibodies. The primary antibody signal was visualized by HRP-conjugated to the corresponding secondary antibodies and the ImageQuant system (GE Imager AI680UV).

For immunoprecipitation with Flag-tagged Cul3, HEK293T cells were homogenized in lysis buffer (50 mM Tris, 200 mM NaCl, 1% NP-40, pH 7.5) with protease inhibitors (Biomake, B14001) for 30 min at 4°C. After remove debris, 20 μl of anti-Flag magnetic beads (Bimake, B26101) were added and the mixture was incubated on a rotator at 4°C overnight. The beads were precipitated using a magnetic stand and washed with cold lysis buffer for 5 times and were resuspend in 3x Protein Loading Buffer and boiling at 95°C for 10 min. The beads were centrifuged at 12,000 rpm for 5 min before SDS–PAGE analysis. The enriched proteins were analyzed by immunoblot (IB).

### Ubiquitination assay

HEK293T cells were transfected with Ub-HA, ZBTB17-Flag and Cul3-V5 plasmids. 36 hours after transfection, cells were treated with 10 μM MG132 (Cayman, 13697-1) for 4 hours. Cells were harvested and were washed 3 times with cold PBS. 100× protease inhibitor cocktail (Biomake, B14001) was added to the lysis buffer prior to the use. Cell pellets were resuspended in 90 μl lysis buffer (20 mM Tris, 150 mM NaCl, 1% Triton X-100, pH7.5) on ice for 1 hour. 10 μl 10% SDS were added to the lysate before sonication. After sonication, the lysate was diluted with lysis buffer to a final concentration of 0.1% SDS. Lysates were centrifuged at 4 °C at 13,000g for 10 min. The supernatant was mixed with 20 μl of Anti-Flag magnetic beads (Bimake, B26101) and incubated on a rotator at 4°C overnight. The beads were washed with cell lysis buffer containing 500 mM NaCl for 5 times and were heated in 3× denaturing loading buffer for 10 min at 95 °C before being resolved by SDS–PAGE.

### RNA extraction and quantitative PCR

500 μl RNAiso Plus (takara, 9109) were added to 0.5 x 10^6^ Cells with immediate mixing. The mixture was then added 100 μl Trichloromethane and inverted 15 seconds. After 10 min incubation, centrifuged at 4 °C at 13,000g for 10 min. Take 200 μl of supernatant to a 1.5mL RNA free tube and then mix with an equal volume of isopropanol. The mixture was incubated for 15 mins and centrifuged at 4 °C at 13,000g for 10 min. Remove the supernatant and added 1 mL of 70% ethanol and centrifuge at 13,000g for 10 min at 4 °C. Dry pellets were resuspended in RNA-free water and the RNA concentration was adjusted to 100 ng/uL.

RNA was reverse transcribed using the PrimeScript™ RT reagent Kit (Takara, RR037A) and the cDNA was used for qPCR using the TB Green® Premix Ex Taq™ (Takara, RR420A), following the supplied protocol. The PCR was running in a QuantStudio 7 Real-Time PCR system. Samples were normalized to Rpl13a gene expression. Primer sequences used in qRT-PCR are presented in **Supplementary Table 7**.

### RNA-seq and ChIP-seq

For RNA-seq, 3 independent biological replicates of experiments were performed. Cells were sent for RNA-seq (Majorbio company). The library was prepared with Illumina TruseqTM RNA sample prep kit and End Repair Mix adaptor sequences have been used. Sequencing was performed with Illumina HiSeq. Gene expression profile changes between control shNT cells and shZBTB17 cells were analyzed using Gene Cluster Analysis and KEGG pathways that were differentially regulated in the absence of functional peroxisomes.

HeLa wild type cells were prepared for ChIP-seq with GENFUND. Cells were crosslinked with 1 % formaldehyde (sigma, 252549) for 10 min at room temperature and quenched with 125 mM glycine (Sangon Biotech, A610235). The fragmented chromatin fragments were pre-cleared and then immunoprecipitated with Protein A + G Magnetic beads (Milliproe, 16-663) coupled with ZBTB17 antibody. After reverse crosslinking, ChIP and input DNA fragments were end-repaired and A-tailed using the NEBNext End Repair/dA-Tailing Module (NEB, E7442) followed by adaptor ligation with the NEBNext Ultra Ligation Module (NEB, E7445). The DNA libraries were amplified for 15 cycles and sequenced using Illumina NextSeq 500 with single-end 1×75 as the sequencing mode. For data analysis, raw reads were filtered to obtain high-quality clean reads by removing sequencing adapters, short reads (length <50 bp) and low quality reads using Cutadapt (v2.4) (Bolger et al., 2014) and Trimmomatic (v0.35) (Martin, 2011). Then FastQC (v0.11.5) is used to ensure high reads quality. The clean reads were mapped to the human genome (assembly GRCh38) using the Bowtie2 (v2.1.0) (Langmead and Salzberg, 2012) software. Peak detection was performed using the MACS (v2.1.5) (Zhang et al., 2008) peak finding algorithm with 0.005 set as the p-value cutoff. Annotation of peak sites to gene features was performed using the ChIPseeker R package (Yu et al., 2015).

### Luciferase assay

Luciferase reporter vectors were constructed based on the pGL4.10 (Promega, E6651), a vector without enhancer or promoter elements. To produce vectors containing various putative PEX13 promoter motifs, different DNA fragments were inserted at the BglI site upstream of the Firefly luciferase gene. For luciferase assays, 0.8 x 10^4^ HEK293T cells were co-transfected using PEI with plasmids at a ratio of 5 ng Renilla luciferase (pGL4.74), 50 ng pGL4.10 empty vector or pGL4.10-PEX13 reporter, and 50 ng transcription factor expression vector (pEF6a-ZBTB17 or pEF6a empty vector) for 24 hours. Luciferase assays were performed in 96-well plates using the Dual-Luciferase Reporter Assay (Promega, E1960) and results were quantified using luminescence counter (SpectraMax i3). Firefly luciferase activity values were normalized to Renilla luciferase activity.

### Subcellular fractionation

5 x 10^6^ HeLa cells were harvested and washed with PBS. 500 μl Peroxisome Extraction Buffer (5 mM MOPS, pH 7.65, with 0.25 M sucrose, 1 mM EDTA, and 0.1% ethanol, Protease Inhibitor Cocktail) was added to cell pellet to achieve an even suspension. The suspended cells were homogenized in a 2 ml Dounce homogenizer (Sigma, D8938-1SET) using Pestle B. Centrifuge the sample at 800 × g for 15 min at 4 °C. The supernatant was subjected to BCA Protein Assay (TIANGEN, PA115-01) to quantify protein levels, the same amount of proteins from each sample were centrifuged at 2,300 × g for 15 min at 4 °C to obtain the pellet. Transfer the supernatant to a new centrifugation tube, centrifuge at 23,000 × g for 20 min at 4 °C. After centrifugation, the pellet (peroxisome fraction) and supernatant (cytosolic fraction) were harvested for further analysis.

### Catalase activity assay

Catalase activity was conducted with peroxisome and cytosolic fraction respectively by using the CheKine^TM^ Catalase (CAT) Activity Assay Kit (Abbkine, KTB1040). 20 μl samples from above were subjected to catalase activity assay as described in the instruction by measuring A_540_ using a SpectraMax i3 microplate Reader.

### Metabolites extraction and metabolomics

HeLa cell samples (shZBTB17, shPEX13 and shNT) were placed in a 6 cm dish and extracted using a metabolite extraction solution MeOH: ACN: H_2_O (2:2:1, v/v) solvent mixture. Place the dishes on dry ice and added 1 mL of cold solvent to each cell plate, and incubate the plates in -80℃ for 40 min. The samples were then scraped and transferred to a 1.5mL EP tube, vortexed for 1 mine and sonicated for 10 min in ice-bath. To precipitate proteins, samples were centrifuged at 13,000 rpm for 10 min at 4 °C. The resulting supernatant was taken and evaporated to dryness in a vacuum concentrator. The dry extracts were then reconstituted in 100 μL of ACN: H_2_O (1:1, v/v), sonicated for 10 min, and centrifuged at 13,000 rpm for 15 min at 4 °C to remove insoluble debris. The supernatant was then transferred to HPLC vials and kept at −80 °C until LC–MS analysis. For each condition, 6 biological repeats were used. For metabolomics, samples were acquired using a UHPLC system (Vanquish, Thermo Scientific) coupled to an orbitrap mass spectrometer (Exploris 480, Thermo Scientific). A Waters BEH amide column was used for LC separation. Mobile phases, linear gradient eluted and ESI source parameters followed the previous publication (cite MetTracer paper). Metabolite annotation was performed using MetDNA (http://metdna.zhulab.cn/). (Shen et al., 2019a; Wang et al., 2022; Zhou et al., 2022a) Metabolomics data was normalized to sample protein concentrations. Metabolic pathway-enrichment analysis was performed via hypergeometric test and visualized in R (v 4.3.0). The pathway database was KEGG (https://www.genome.jp/kegg/).

## Data Availability

The RNA sequencing data and ChIP-sequencing data had been deposited in the NCBI Gene Expression Ominibus (GEO) under the accession number GSE239792.

## ACKNOWLEDGMENTS

We would like to thank the Molecular Imaging Core Facility (MICF), and the Molecular and Cell Biology Core Facility (MCBCF) in the School of Life Science and Technology at ShanghaiTech University for providing technical support. M.Z. is supported by the National Key R&D Program of China (2021YFA0804700 and 2021YFA1100800), Shanghai Frontiers Science Center for Biomacromolecules and Precision Medicine at ShanghaiTech University, and the Innovative Research Team of High-level Local Universities in Shanghai.

The authors declare no competing financial interests.

---

**Figure S1.**
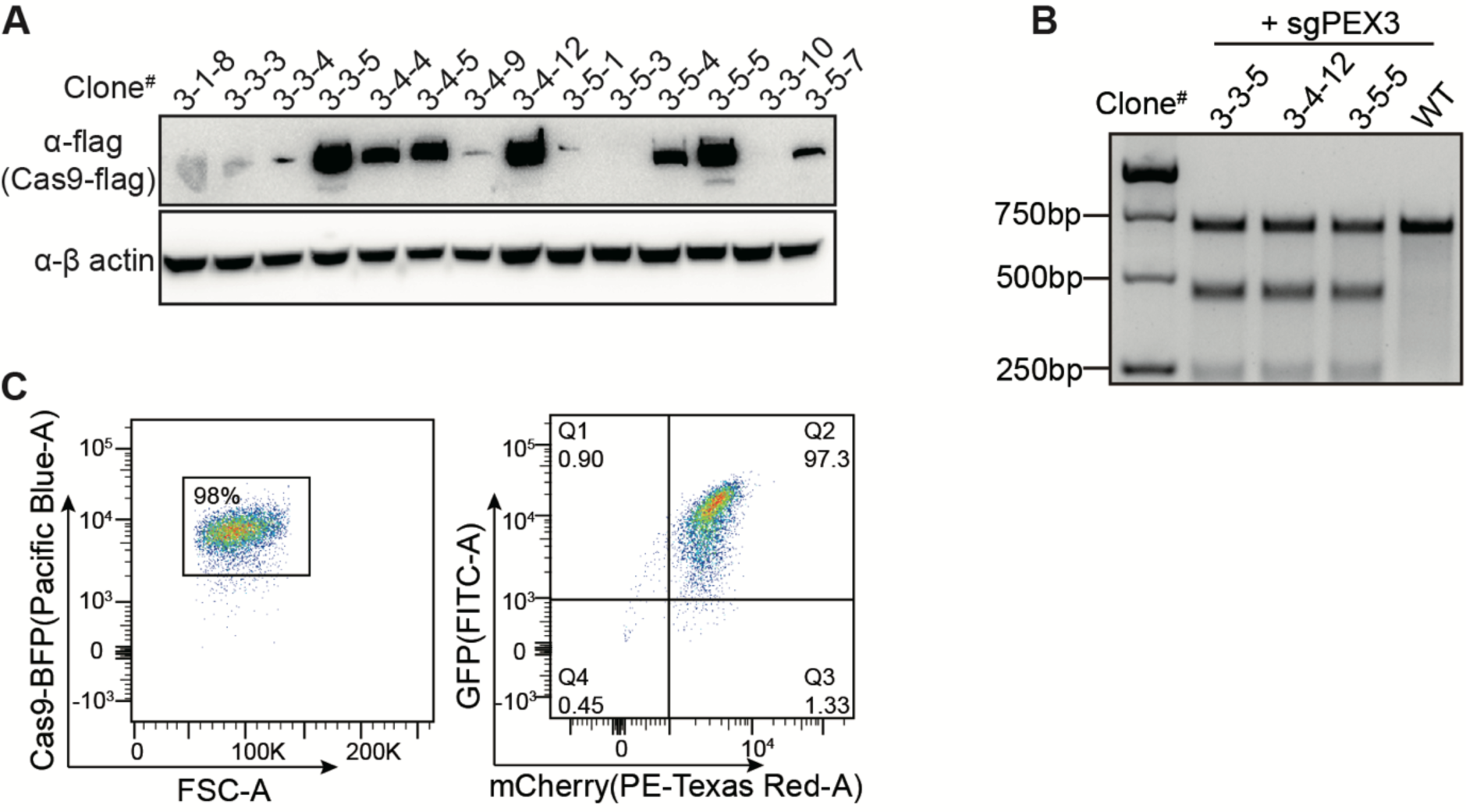
Isolation of a single cell clone containing both Cas9 and the reporter. **(A)** Cas9 expression level detected with anti-flag antibody in different clones. **(B)** Lentivirus-delivered sgRNA targeting the PEX3 gene in the indicated cell clones were assayed by T7E1 digestion, showing Cas9 is active in the stable cell line. **(C)** Generation of Cas9 and EGFP/mCherry containing cell clone.

**Figure S2.**
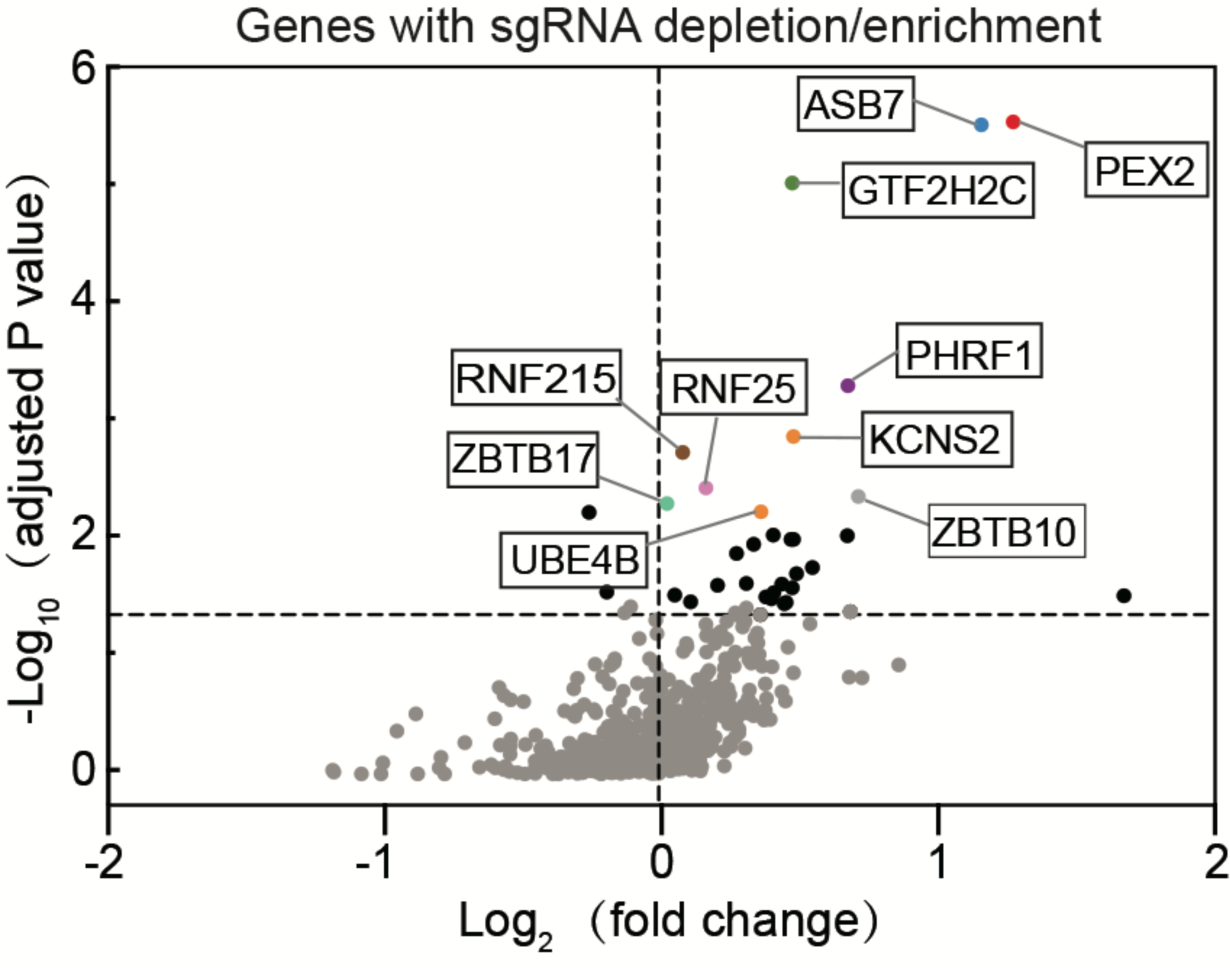
Volcano plot illustrating the differential expression of sgRNAs (shown as the name of the targeting genes). The fold change indicates the enrichment of these sgRNAs in the lowest 30% of the GFP/mCherry cells compared to highest 30% of the GFP/mCherry cells. The top candidate genes are highlighted.

**Figure S3.**
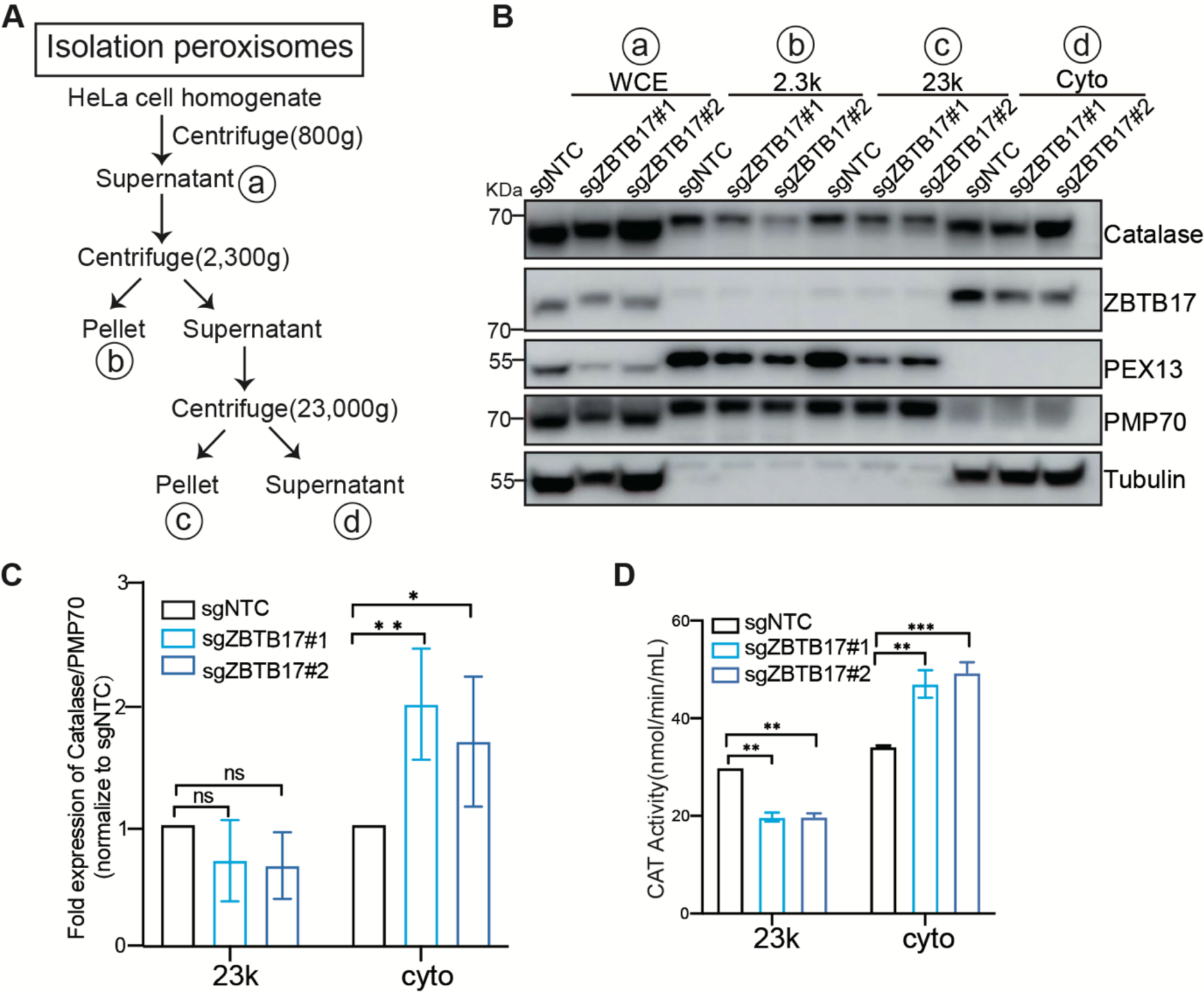
Catalase is redistributed in ZBTB17 deficient cells. **(A)**Flowchart of peroxisome purification procedure. **(B)** Western blot analysis of catalase in different fractions in cells with or without ZBTB17 knockout. Cellular fractionation experiments were performed using stepwise centrifugation, equal volume of samples with each fraction isolated from cells was analyzed with IB using the indicated antibodies. WCE: whole-cell extraction; 2.3K: pellet after centrifugation at 2,300 g; 23K: the major peroxisome fraction, pellet after centrifugation at 23,000 g; Cyto: supernatants after centrifugation at 23,000 g. **(C)** Catalase levels in various fractions of distinct samples (F) were quantified and normalized using ImageJ. Values are mean ± SD, n = 3 independent experiments. *, P < 0.05; ***, P < 0.001; ****, P < 0.0001; n.s., not significant (P > 0.05) by two-way ANOVA multiple comparisons test. **(D)** Catalase activity in various fractions of distinct samples (F) were measured. Values are mean ± SD, *, P < 0.05; ***, P < 0.001; ****, P < 0.0001; n.s., not significant (P > 0.05) by two-way ANOVA multiple comparisons test (n=2).

**Figure S4.**
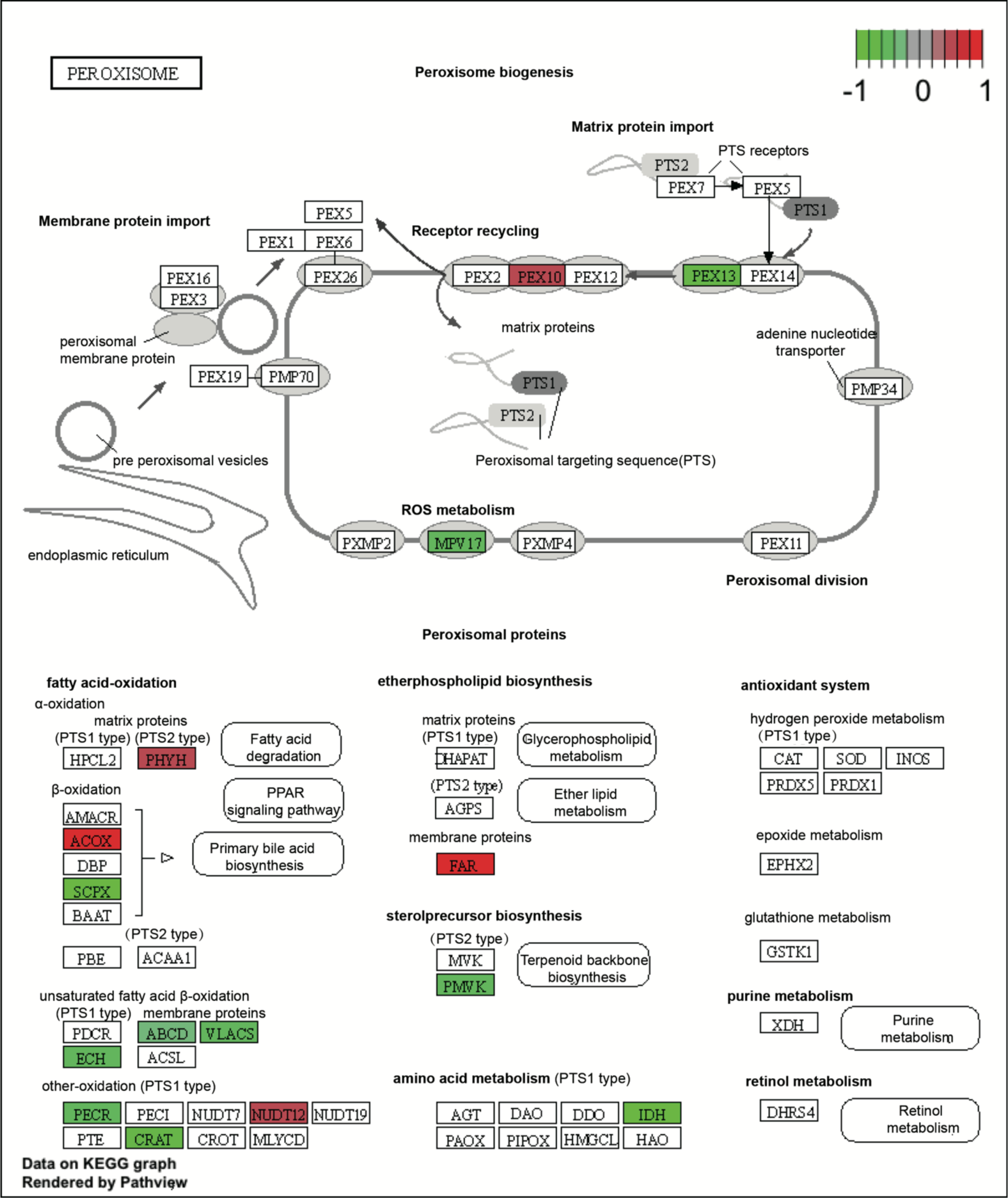
Peroxisome-related genes with altered expression highlighted in KEGG (Kyoto Encyclopedia of Genes and Genomes) pathway. Red, up-regulated genes; green, down-regulated genes. This visualization was conducted using the R package ‘pathview’ 1 on KEGG graphs (p.adjust < 0.05).

**Figure S5.**
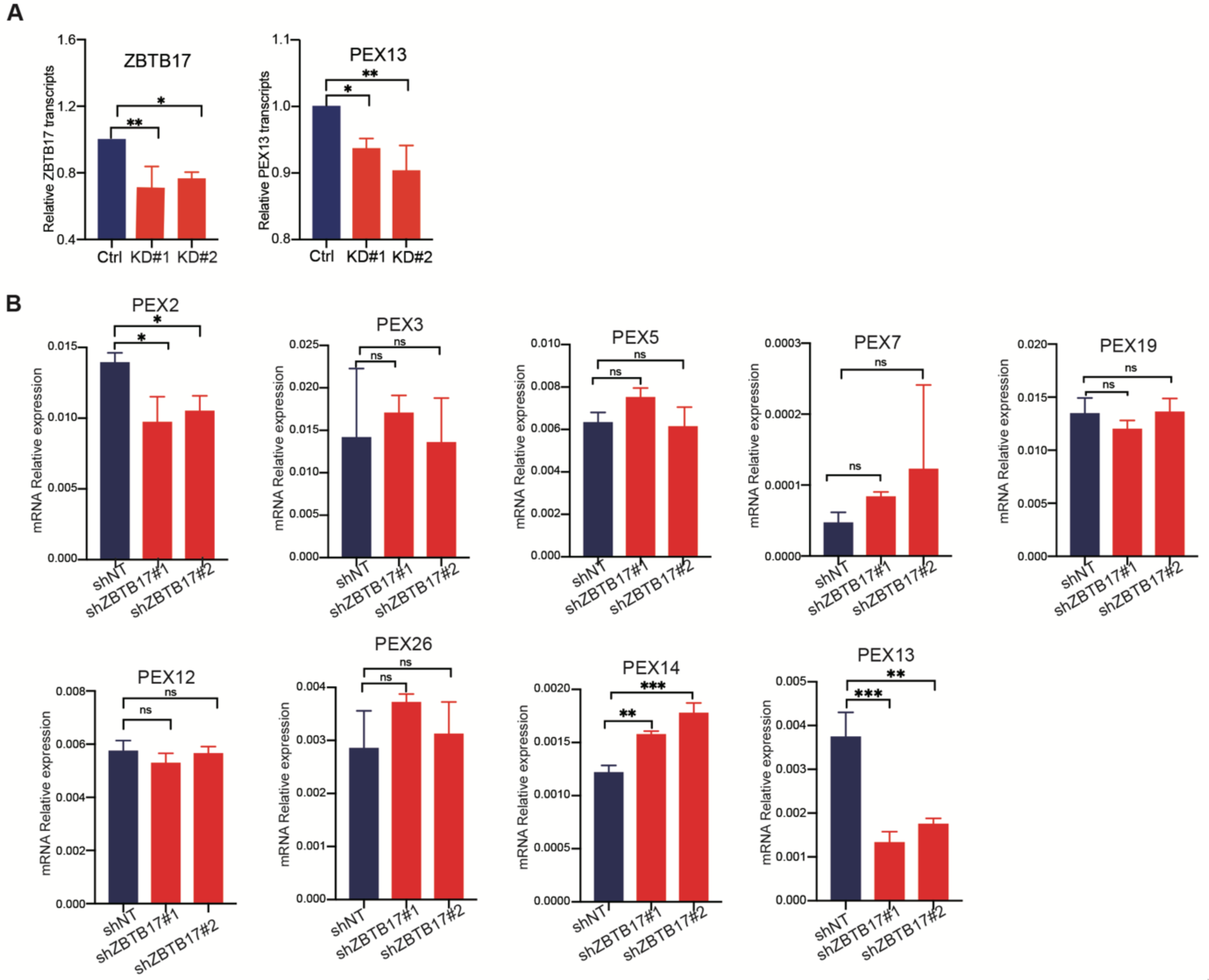
mRNA levels of different peroxin genes. **(A)** Relative levels of ZBTB17 and PEX13 transcripts in ZBTB17 knockdown and wild-type HeLa cells, as determined from RNA-seq data. **(B)** Quantitative PCR (Q-PCR) analysis of the mRNA levels of peroxisome transport genes in ZBTB17 knockdown and wild-type HeLa cells. Statistically significant differences are indicated (P<0.05*, P<0.01**, P<0.0001***).

**Figure S6.**
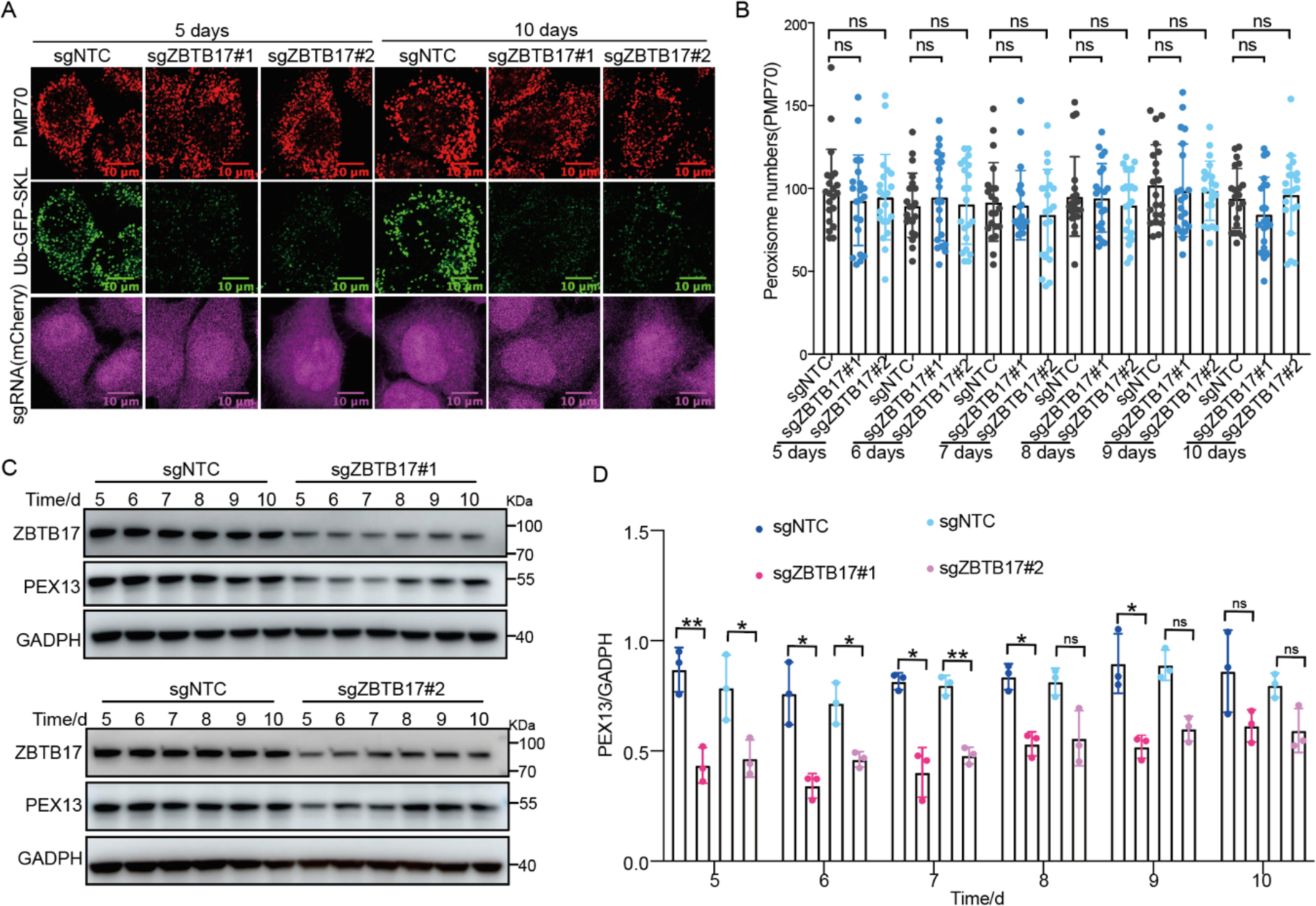
Peroxisome numbers do not change over time. **(A)** Ub-GFP-SKL HeLa cells were transfected with sgZBTB17 for 10 days and stained for PMP70. Representative immunofluorescence microscopy images of 5-day and 10-day are shown. **(B)** Quantification of peroxisome numbers (PMP70 puncta) in > 20 cells in 5 or 10 days. **(C)** Immunoblotting of PEX13 and ZBTB17 in HeLa cells transfected with sgRNA/Cas9 targeting ZBTB17 for 5 to 10 days. (D)PEX13 levels in (C) were quantified and normalized using ImageJ. Values are mean ± SD, n = 3 independent experiments. *, P < 0.05; ***, P < 0.001; ****, P < 0.0001; n.s., not significant (P > 0.05) by two-way ANOVA multiple comparisons test.

## Supplementary Tables

Table S1 The sgRNA sequences used in this study.

Table S2 The E3 library of sgRNA sequence. (<u>as a separate file</u>)

Table S3 The complete list of sgRNAs ranking with CRISPR screen, related to **Figure 1D, Figure S2**. (<u>as a separate file</u>)

Table S4. Differentially expressed genes in ZBTB17 KD cells. (<u>as a separate file</u>)

Table S5 Metabolomic identifications (as a separate file)

Table S6 The shRNA sequences used in the study

Table S7 Summary of primer sequences for quantitative PCR.

**Supplementary Table S1.**
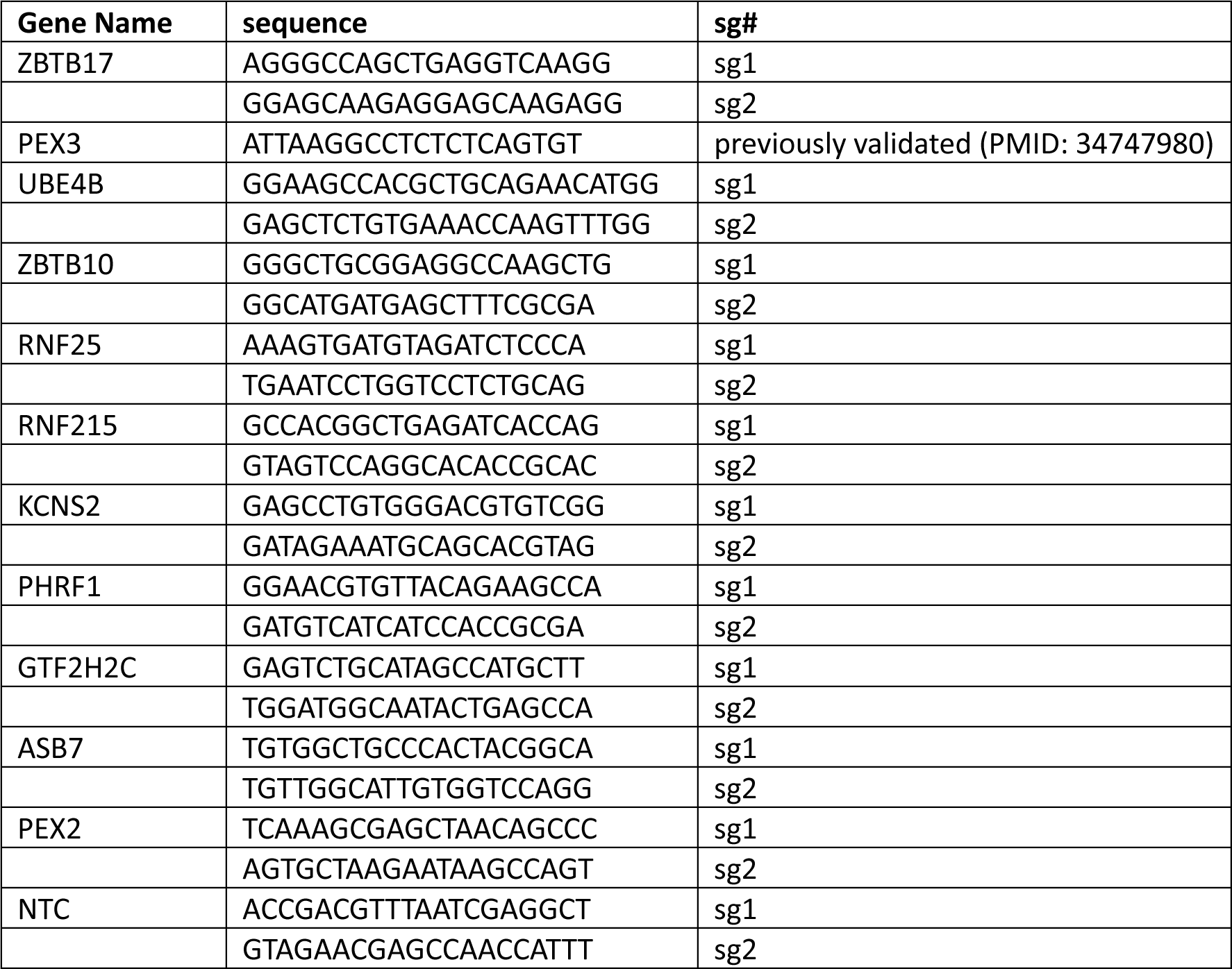
The sgRNA sequences used in this study.

**Supplementary Table S6.**
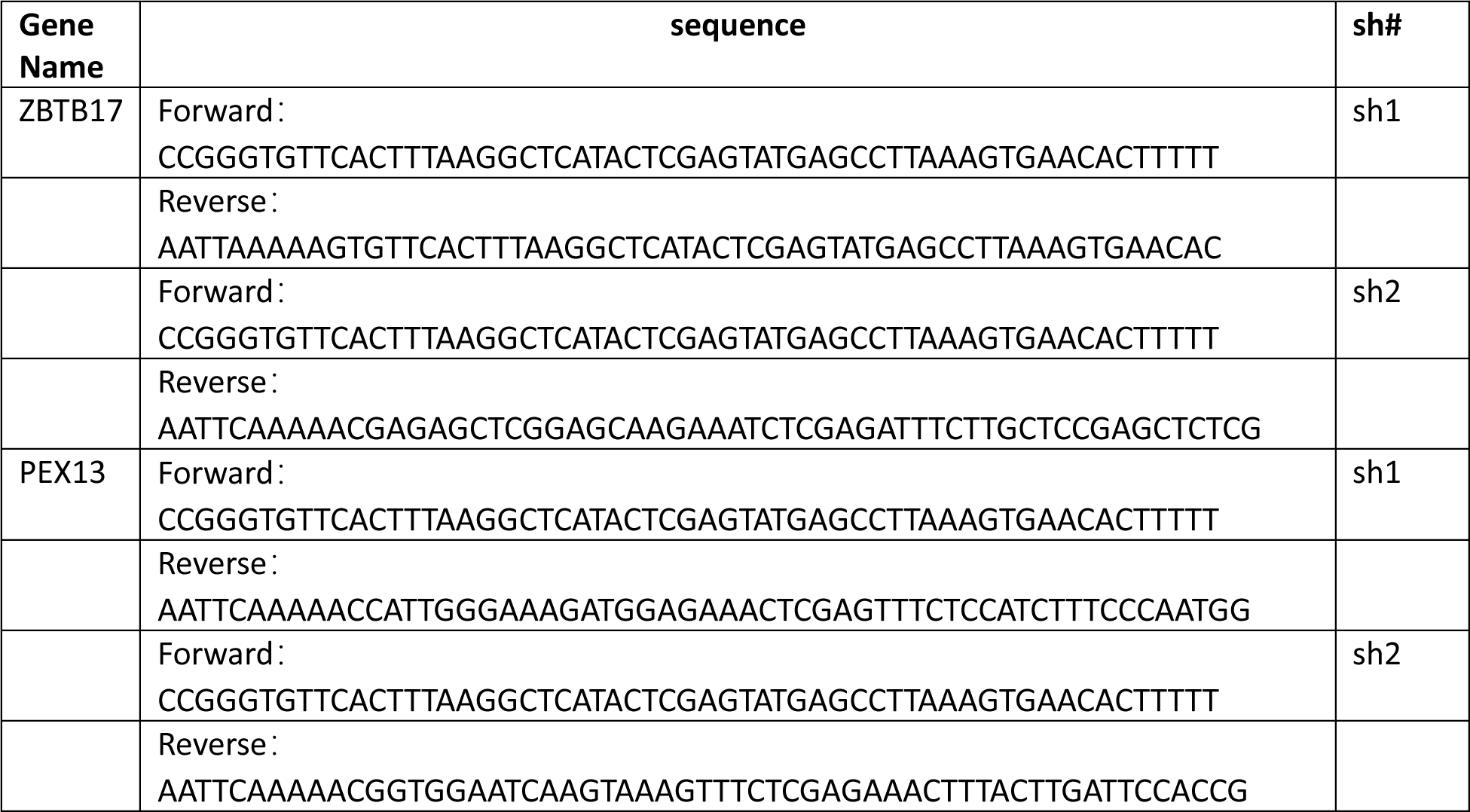
The shRNA sequences used in this study.

**Supplementary Table 7.**
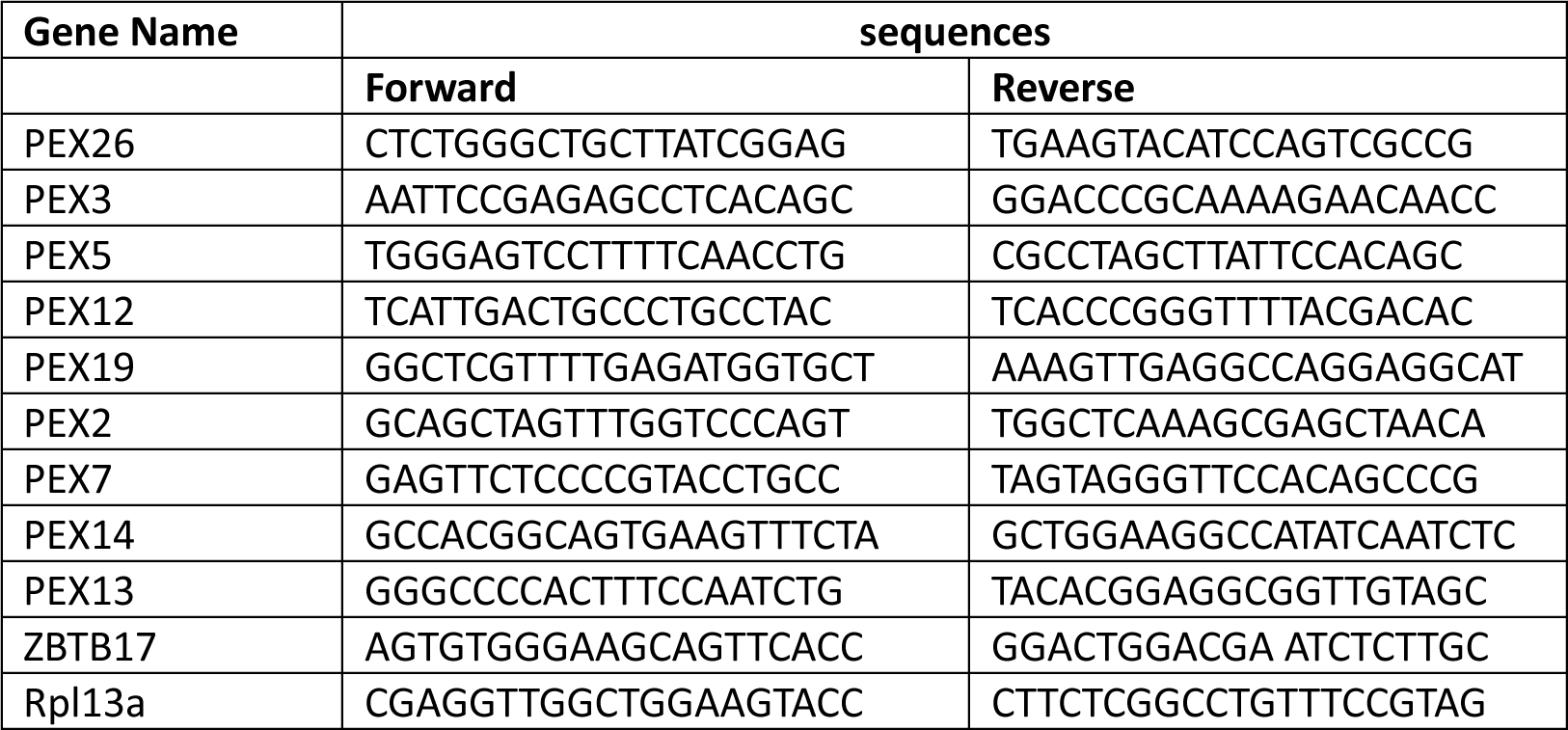
Summary of primer sequences for quantitative PCR.

